# N^6^-Methyladenosine Safeguards Mouse and Human Germline Competence

**DOI:** 10.1101/2025.11.18.688769

**Authors:** Rujuan Zuo, Baoyan Bai, Kang-Xuan Jin, Mirra Louise Cicilie Søegaard, Erkut Ilaslan, Ying Yao, Jingwei Li, Łukasz Wyrożemski, Yanjiao Li, Xuechen Wu, Junbai Wang, Arne Klungland, Mads Lerdrup, Adam Filipczyk

**Affiliations:** Centre for Embryology and Healthy Development, Division of Laboratory Medicine, Laboratory for Stem Cell Dynamics and RNA Regulation, Oslo University Hospital, Forskningsveien 1, SINTEF Building 0373, Oslo; Centre for Embryology and Healthy Development, Division of Laboratory Medicine, Laboratory for Dynamic Gene Regulation, University of Oslo, and Oslo University Hospital, Forskningsveien 1, SINTEF Building 0373, Oslo; Clinical Molecular Biology (EpiGen), Medical Division, Akershus University Hospital and Institute of Clinical Medicine, University of Oslo, Sykehusveien 25, 1447, Lørenskog; Department of Cellular and Molecular Medicine, The DNRF center for Chromosome Stability, University of Copenhagen, Blegdamsvej 3, København; Department of Immunology, University of Oslo, Problemveien 7, 0315 Oslo

**Keywords:** N^6^-Methyladenosine, m^6^A, formative pluripotency, germ cell, epigenetics, signaling

## Abstract

N^6^-Methyladenosine (m^6^A) regulation of germline entry in post-implantation-like-pluripotent states remains poorly defined. We characterized m^6^A depletion by *Mettl3* or *Mettl14* knock-down or METTL3 inhibition in *in* vitro models enabling primordial germ-cell-like-cell and somatic differentiation. Upon m^6^A depletion, mouse epiblast-like cells upregulated embryonic *Ras* (Eras) and PI3K-pAKT, facilitating EZH2 repression and depletion of H3K27me3. Consequently, enhancer activation promoted global gene deregulation and impaired germline entry. In mouse formative stem cells, ERAS-PI3K-pAKT upregulation resulted in unscheduled *Blimp1* and *Otx2* co-expression, germline entry deficiency and enhanced mesodermal differentiation. Conversely, upon m^6^A depletion, germline competent human pluripotent stem cells upregulated OTX2 through FGF-pERK activation, resulting in germline entry deficiency and enhanced neuroectodermal differentiation. meRIP-seq revealed preferential m^6^A deposition on mouse mesodermal and human neuroectodermal transcripts, suggesting a basis for differentiation bias following m^6^A depletion. We illustrate how decreasing m^6^A abundances impact signaling, chromatin epigenetics and transcript stabilities, constituting a powerful cell intrinsic barrier for germline entry.

## Introduction

Understanding mechanisms governing the acquisition of competence for lineage entry will enable new strategies to obtain functional tissues for stem cell-based therapies. Naïve mouse embryonic stem cells (mESCs) progress through formative pluripotency to acquire germline competence and the responsiveness to lineage induction signals at gastrulation^1-3^. The formative phase was first emphasized in transient, germ-cell-competent mouse epiblast-like-cells (EpiLCs), which can be propagated for 2 days but cannot be maintained in this state^4^. 5-25% of day 2 EpiLCs achieves sensitivity to Fibroblast growth factor (FGF) and NODAL signaling as well as BMP responsiveness, which is critical for primordial germ cell fate initiation^4^. Mouse germline entry is characterized by the establishment of a tripartite germ cell transcription factors (TFs) network composed of BLIMP1, PRDM14 and AP2γ (encoded by *Tfap2c*) expression^5-7^. Having passed the formative stage, primed epiblast stem cells (EpiSCs) respond poorly to germ cell inductive signals^4,8^. Recent reports show that FGF and addition of Activin alongside inhibition of WNT captures self-renewing formative stem (FS) cells^2^ and formative pluripotent stem cells (fPSCs)^9^. Alternatively, PSCs grown in FGF, TGF-β and WNT agonist supplemented media (FTW-PSCs) harbor early formative features^10^. As in EpiLCs, these conditions promote the downregulation of naïve TFs like Nanog and *Klf4* and the establishment of a formative TF network incorporating *Otx2* and *Oct6*. Lineage TFs found in mEpiSCs, like *T* and *Sox1* remain downregulated^2,10^.

By contrast, early human post-implantation epiblast retains persistent *NANOG* and *PRDM14* expression alongside low *OTX2* levels^11,12^. Developmental progression involves *SOX11* upregulation^11,12^ and on the verge of gastrulation, the emergence of lineage TFs like *T, GATA4, GATA6*, and *MIXL1*^3,11^. Using conditions similar to mouse FS cell culture, a handful of human formative stem cell populations have been expanded, although cell state heterogeneity and germline competence in these systems require further investigation^2,10^. Cells adapted from conventional human pluripotent stem cell (hPSC) culture, also retain properties of the formative landscape^3,13-15^. For example, hPSCs adapted to 4i conditions, supplemented with GSK3, ERK, JNK and P38 inhibitors, bFGF, TGF-β and LIF (Leukemia Inhibitory Factor), show robust germline competence^14,16^. A recent comparative gene expression analysis places 4i hPSCs alongside the human early post-implantation epiblast (embryonic day 8) and FTW-PSCs^10^. 4i hPSCs were used to characterize *SOX17* as a central regulator of human primordial germ cell-like cell (hPGCLC) lineage entry and *BLIMP1* as a downstream repressor of somatic lineage induction^14^. Other cells derived from hPSCs and capable of germline entry are a transient population termed induced mesodermal like cells (iMeLCs)^13^. They are generated in Activin A and WNT agonist CHIR99201 supplemented media, although unlike 4i hPSCs, bFGF supplementation has proven detrimental to iMeLC expansion^13^. Either method likely induces a heterogeneous mix of poorly defined peri-gastrulating epiblast-like cells, which include a subpopulation capable of germline competence acquisition^3^. Upon BMP4 addition, this fraction can range from approximately 5-35%, depending on the induction method and cell background^13^.

Recently, epi-transcriptomic regulation has emerged as central to developmental progression^17^, with open questions about how a dynamic epi-transcriptome governs formative pluripotency in human and mouse. N^6^-Methyladenosine (m^6^A) is the most abundant messenger RNA (mRNA) modification^18^, present in at least a quarter of all mRNAs and typically near stop codons^19^. A multi-protein ‘writer’ complex deposits m^6^A and includes the methyltransferase METTL3, tasked with m^6^A deposition. The reversibility of m^6^A by two demethylases or ‘erasers’ (ALKBH5 and FTO) suggests dynamic cell fate regulation^17,20-22^. Reader proteins like YTHDFs or IGFBPs recognize m^6^A and coordinate transcript stability, splicing, or translation^17,22,23^. Interestingly, both *Nanog* and *Otx2* transcripts are co-expressed in mouse and human formative cells^2,10^. The transcript stability of either TF may be elevated upon m^6^A depletion^24,25^. In mouse EpiLCs, increased *Nanog* expression alone is sufficient to induce mouse primordial germ-cell-like-cells (mPGCLCs)^26^, while *Otx2* is a powerful germline repressor^27^. Recently in transient human iMeLCs, *OTX2* transcript stability was shown to be increased by IGF2BP1 binding in an m^6^A dependent manner^28^. Hence, upon m^6^A depletion iMeLC germline entry was increased^28^. However, it remains unclear whether decreasing m^6^A abundances are conducive to germline competence in other formative cell populations, including mouse (EpiLCs and FS cells) or self-renewing human germline competent cells, like 4i hPSCs. Furthermore, how formative cells preclude unscheduled germline entry amid fluctuating m^6^A levels remains open to debate^17^.

In this study, we show that m^6^A abundances safeguard efficient germline entry in mouse and human formative cells. Depletion of m^6^A in EpiLCs elevated ERAS transcript stability, fostering ERAS-PI3K-pAKT signaling activation and facilitating EZH2 downregulation. Consequently, global remodeling of low abundance H3K27 trimethylation found on enhancers, was associated with gene deregulation and germline entry deficiency. Depletion of m^6^A in FS cells showed unscheduled co-expression of *Otx2* and mesendodermal genes together with transcript stability dependent upregulation of *Eras* and *Blimp1*. Germline entry deficiency was accompanied by mesodermal differentiation partiality. By contrast in human 4i hPSCs, m^6^A depletion activated FGF-pERK signaling, promoting increased OTX2 expression. Restricted germline competence was accompanied by neuroectodermal gene upregulation and increased propensity for neuroectodermal differentiation. meRIP-seq analysis showed preferential m^6^A deposition on mouse mesodermal and human neuroectodermal transcripts, potentially underlying differentiation biases observed upon m^6^A depletion. In light of recent work^28^ and models showing m^6^A abundances regulating developmental progression^17^, we illustrate how decreasing m^6^A abundances establishes a powerful cell intrinsic barrier to germline entry.

## Results

### Depletion of m^6^A Impairs Germline Entry

We depleted m^6^A in naïve 2i+LIF mESCs by *Mettl3* or its methyltransferase complex partner *Mettl14* shRNA mediated knock-down. Upon m^6^A depletion, cells were expanded for 10-days before initiating 2-days of EpiLC induction, with no noticeable change to EpiLC morphology or growth (Supplementary Fig. 1a). RT-qPCR analysis for *Mettl3* and *Mettl14* mRNA levels confirmed that *Mettl3* and *Mettl14* knock-down in naïve mESCs persisted in day 2 EpiLCs and 4 days after mPGCLC induction (Fig. 1a). Knock-down of *Mettl3* or *Mettl14* corresponded with METTL3 and METTL14 protein down-regulation (Supplementary Fig. 1b). This significantly reduced global m^6^A abundances as shown by dot-blot profiling (Supplementary Fig. 1c) and mass spectrometry analysis (Supplementary Fig. 1d). In contrast, no changes were found by analysis of m^7^G or m^6^Am abundances (Supplementary Fig. 1d). m^6^A depleted day 4 mPGCLC spheres retained significantly lower expression of *Blimp1, Prdm14, Nanos3, Stella, Tfap2c*, and *Dnd1* germ cell genes, indicative of germline entry deficiency (Fig. 1a). CD61 and SSEA1 surface marker double positive cells also account for mPGCLCs expressing *Blimp1* and *Stella*^4^. Depletion of m^6^A significantly reduced the CD61 and SSEA1 double positive cell fraction (Fig. 1b, c). This corresponded with a decrease in the proportion of *Blimp1*-GFP fluorescence reporter positive cells (Supplementary Fig. 1e). Depletion of m^6^A and reduction in germline entry, was further recapitulated using the analogous fPSC system (Supplementary Fig. 1f).

**Figure 1.**
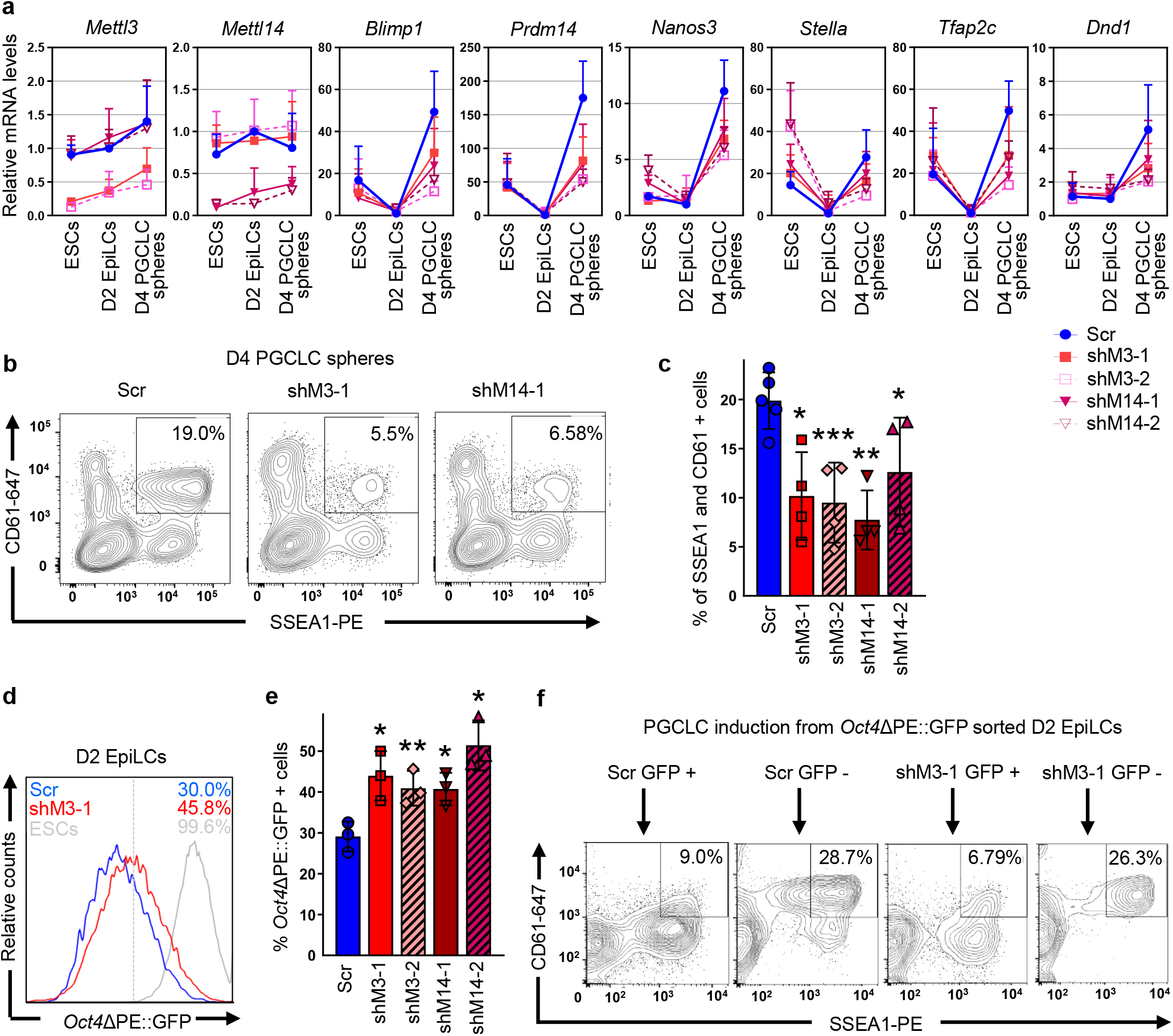
Depletion of m^6^A Impairs Germline Entry. **a-e**, *Oct4*ΔPE::GFP or *Blimp1*-GFP mESCs were transduced with scramble control (Scr), *Mettl3* (shM3-1 or shM3-2) or *Mettl14* (shM14-1 or shM14-2) shRNA, followed by EpiLC induction and PGCLC specification. Data is presented as the mean ± standard deviation (SD). Two-tailed, paired t-test: vs Scr, \*\*\**p* <0.001; \*\**p* <0.01; \**p* <0.05. **a**, qPCR analysis showing shRNA knock-down efficiency and effects on germline markers in the *Blimp1*-GFP cell line. The value for Scr Day 2 EpiLCs was set to 1. N ≥3. Statistical analyses are listed in Supplementary Table 2. **b**, FACS analysis of surface markers SSEA1 and CD61 in *Blimp1*-GFP day 4 PGCLC spheres. Percentages of SSEA1 and CD61 double positive cells are shown. Cells are gated according to SSEA1 and CD61 signals in no cytokine treated control spheres. **c**, Graph summarizing effects in (*b*). N ≥4. **d**, FACS analysis of *Oct4*ΔPE::GFP expression in day 2 EpiLCs. Percentages of *Oct4*ΔPE::GFP positive cells are shown. Dashed line indicates positive gating. Solid gray line represents GFP signals in positive control. **e**, Graph summarizing effects in (**d**). N =3. **f**, *Oct4*ΔPE::GFP mESCs were transduced with scramble control (Scr) or *Mettl3* (shM3-1) shRNA before EpiLC induction. Day 2 EpiLCs were sorted into GFP positive and negative cell fractions for PGCLC specification. FACS analysis measuring percentages of SSEA1 and CD61 double positive cells in day 4 PGCLC spheres. See also Figure S1r for experimental repeats.

The acquisition of competence for germline entry in day 2 EpiLCs is associated with epigenetic enhancer remodeling^4,26,29-31^. At the *Oct4* locus, this process is represented by the switching of enhancer usage from distal to proximal. This was previously shown by the loss of *Oct4*ΔPE::GFP fluorescence reporter which delineates *Oct4* distal enhancer activity^26,32-34^. Depletion of m^6^A significantly increased the *Oct4*ΔPE::GFP positive cell fraction in day 2 EpiLCs and the effect was inversely proportional to the reduction in germline entry (Fig. 1d, e). We next asked if *Oct4*ΔPE::GFP retention in m^6^A depleted day 2 EpiLCs was a by-product of hyper-pluripotency, reported upon stable m^6^A depletion in naïve mESCs^25,35^. Hyper-pluripotency is characterized by persistent *Nanog* expression and consequently, poor capacity to exit naïve pluripotency^25,35^. To exclude this possibility, acute depletion of m^6^A in mESCs showed robust down-regulation of the *Nanog*VENUS protein-fusion reporter^36,37^ (Supplementary Fig. 1g) and total NANOG protein levels in EpiLCs (Supplementary Fig. 1h). Additionally, cells at day 1 of EpiLC induction were previously shown to re-acquire naïve pluripotency and re-instate *Oct4*ΔPE::GFP expression when re-introduced into 2i+LIF. However, by day 2 of induction, EpiLCs permanently lost this capacity^26^. In 2i+LIF, we found that a lower proportion of m^6^A depleted day 1 EpiLCs retained *Oct4*ΔPE::GFP (Supplementary Fig. 1i) and by day 2, cells lost the ability to re-express *Oct4*ΔPE::GFP altogether (Supplementary Fig. 1i). Finally, m^6^A depletion was achieved in EpiLCs by *Mettl3* or *Mettl14* knock-down (Supplementary Fig. 1j) or METTL3 catalytic inhibitor (STM2457) treatment^38^. STM2457 significantly reduced total m^6^A levels as confirmed by ELISA assay (Supplementary Fig. 1k) and mass spectrometry analysis (Supplementary Fig. 1l). Either *Mettl3* or *Mettl14* knock-down (Supplementary Fig. 1m) or STM2457 (Supplementary Fig. 1n), increased the proportion of *Oct4*ΔPE::GFP positive EpiLCs. In line with these observations, STM2457 treatment resulted in impaired germline entry as measured by the CD61 and SSEA1 double positive cell fraction (Supplementary Fig. 1o).

To confirm that *Oct4*ΔPE::GFP persistence decreased germline competence in day 2 EpiLCs, control and m^6^A depleted cells were fluorescence sorted into GFP positive and negative cell fractions (Supplementary Fig. 1p). In either control or m^6^A depleted condition, GFP negative cells showed efficient mPGCLC induction in contrast to GFP positive cells, where the capacity for germline entry was significantly reduced (Fig. 1f, Supplementary Fig. 1q, r). Therefore, m^6^A depletion sustains *Oct4*ΔPE::GFP in day 2 EpiLCs and represses *Oct4* distal enhancer silencing required for germline competence acquisition.

### m^6^A Dependent Eras Transcript Degradation in EpiLCs Ensures Robust Germline Entry

Genome-wide transcriptome profiling of day 2 EpiLCs identified 12 genes with significantly changed expression and log_2_ fold changes higher than 1 upon either *Mettl3* or *Mettl14* knock-down (adjusted *p* ≤0.05) (Fig. 2a, b). Deviations in the levels of many genes were specific to either *Mettl3* or *Mettl14* depletion (Fig. 2a). This may reflect methyltransferase complex independent roles, recently ascribed to METTL3 or METTL14^39-42^. These candidates were excluded from further study. Pluripotency TFs were not significantly changed in m^6^A depleted day 2 EpiLCs (Supplementary Fig. 2a, b). The germline repressor *Otx2*^27^ was previously shown to undergo upregulation upon m^6^A depletion^24^. However, mRNA expression of *Otx2, Oct6, Fgf5* or other developmental regulators (Supplementary Fig. 2a, b) as well as OTX2 protein levels remained unchanged in m^6^A depleted day 2 EpiLCs (Supplementary Fig. 2c).

**Figure 2.**
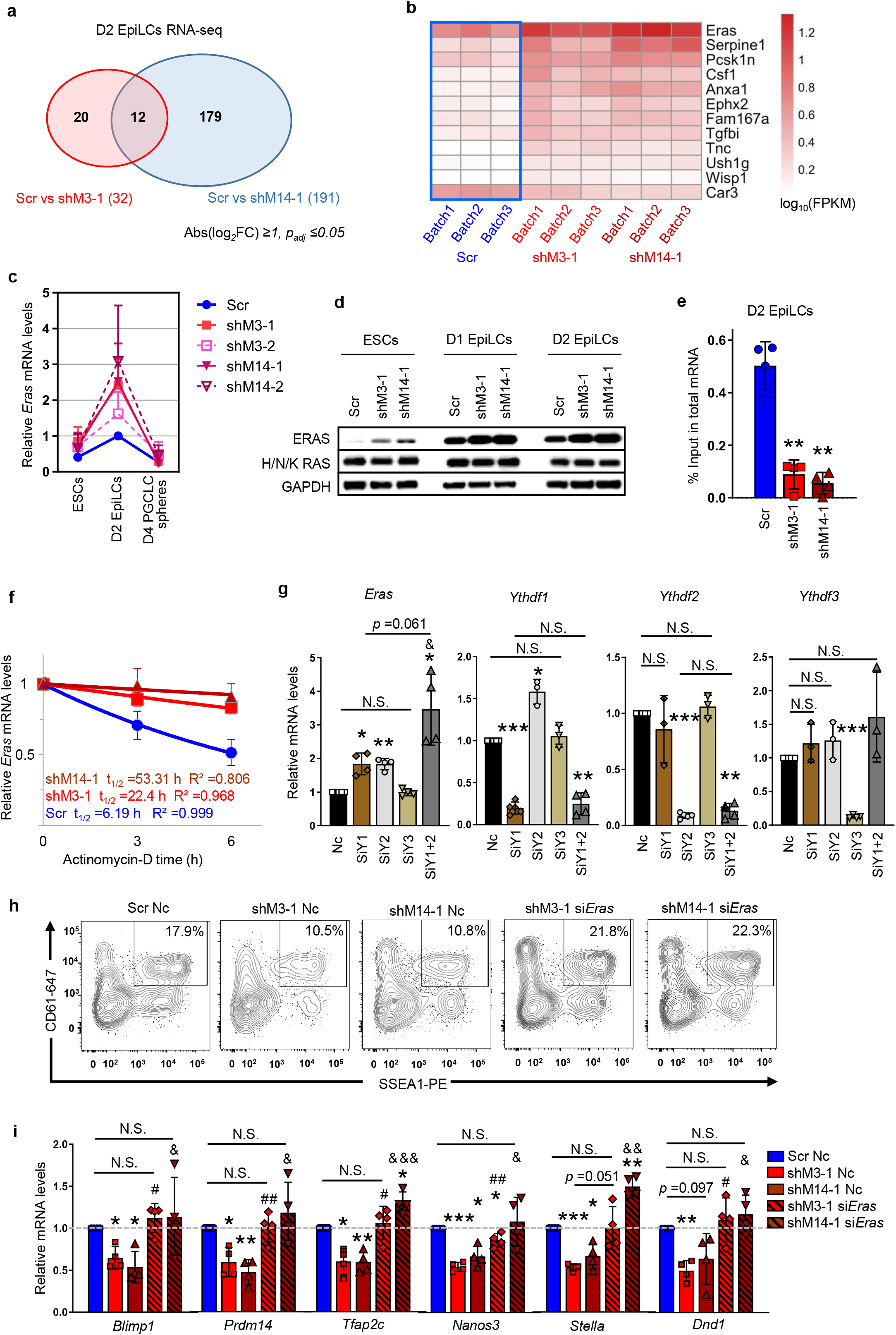
m^6^A Dependent *Eras Transcript Degradation in EpiLCs Ensures Robust Germline Entry*. **a-f**, *Oct4*ΔPE::GFP or *Blimp1*-GFP mESCs were transduced with scramble control (Scr), *Mettl3* (shM3-1 or shM3-2) or *Mettl14* (shM14-1 or shM14-2) shRNA, followed by EpiLC induction and PGCLC specification. **a**, RNA-seq Venn diagram showing overlap of differentially expressed genes (DEGs) in *Oct4*ΔPE::GFP day 2 EpiLCs. The number of DEGs for each condition is shown. Abs(log_2_FC), absolute log_2_Fold Change. *p*_*adj*_, adjusted *p* value. **b**, Heat map showing the expression of 12 DEGs in (a), including three independent experimental repeats. FPKM, Fragments Per Kilobase of transcript per Million mapped reads. **c**, qPCR of *Eras* expression during *Blimp1*-GFP EpiLC and PGCLC inductions. Values were normalized to Scr day 2 EpiLCs defined as 1. Mean ± SD. N ≥5. Statistical analyses are listed in Supplementary Table 2. **d**, Immunoblots showing protein levels of ERAS and H/N/K RAS isoforms in *Blimp1*-GFP ESCs, day 1 EpiLCs and day 2 EpiLCs. **e**, meRIP-qPCR showing m^6^A levels on *Eras* mRNA in *Blimp1*-GFP ESCs and day 2 EpiLCs. Mean ± SD. N =4; two-tailed, paired t-test: vs Scr, \*\**p* <0.01; \**p* <0.05. **f**, qPCR showing *Eras* mRNA half-life (t_1/2_) after Actinomycin-D treatment in *Oct4*ΔPE::GFP day 2 EpiLCs. Mean ± SD. N =3. Statistical analyses are listed in Supplementary Table 2. **g**, O*ct4*ΔPE::GFP day 0 EpiLCs were transfected with negative control (Nc), *Ythdf1* (siY1), *Ythdf2* (siY2), *Ythdf3* (siY3) or *Ythdf1* and *Ythdf2* (siY1+2) siRNA. qPCR in day 2 EpiLCs showing *Eras* expression and knock-down efficiency (N ≥4), Mean ± SD. Two-tailed, paired t-test: vs Nc, \*\*\**p* <0.001; \*\**p* <0.01; \**p* <0.05; vs siY2,^&^*p* <0.05; N.S., Non-significant. **h-i**, *Blimp1*-GFP reporter ESCs were transduced with scramble control (Scr), *Mettl3* (shM3-1) or *Mettl14* (shM14-1) shRNA before EpiLC induction. Day 0 EpiLCs were transfected with negative control (Nc) or *Eras* (si*Eras*) siRNA for 2 days, followed by PGCLC specification. See also Figure S2m for experimental details. **h**, FACS analysis showing percentages of SSEA1 and CD61 double positive cells in day 4 PGCLC spheres. See also Figure S2p for experimental repeats. **i**, qPCR of germline markers in day 4 PGCLC spheres. Mean ± SD. N =4; two-tailed, paired t-test: vs Scr Nc, \*\*\**p* <0.001; \*\**p* <0.01; \**p* <0.05; vs shM3-1 Nc, ^##^*p*<0.01; ^#^*p* <0.05; vs shM14-1 Nc, ^&&&^p <0.001; ^&&^*p* <0.01; ^*^*p* <0.05; N.S., non-significant.

We focussed on *Eras* for further investigation as it encodes a constitutively active Ras-like GTPase protein that stimulates phosphatidylinositol-3-OH kinase (PI3K) dependent pAKT signaling^43^. PI3K-pAKT is activated downstream of FGF signaling during epiblast differentiation^44^. *Eras* was originally identified in mESCs where it facilitates cell growth and teratoma forming properties^43,45^. However, the expression of *Eras* persists in the peri-and post-implantation mouse epiblast^46^. Its role in formative pluripotency remains unexplored. RT-qPCR analysis confirmed that the expression of *Eras* was highest in m^6^A depleted day 2 EpiLCs (*p* <0.05, two-tailed, paired t-test) relative to levels in mESCs and mPGCLC spheres (Fig. 2c). This was also reflected by ERAS protein levels in EpiLCs compared to mESCs, while other *Ras* oncogene isoforms remained unchanged (Fig. 2d). Sequence analysis predicted that *Eras* mRNA contains multiple putative m^6^A motif sites (Supplementary Fig. 2d), and can be m^6^A modified^25^. Methylated-mRNA-immunoprecipitation-qPCR (meRIP-qPCR) confirmed that the 3’ end of *Eras* mRNA coding sequence (CDS) region undergoes a significant reduction in m^6^A deposition upon *Mettl3 or Mettl14* knock-down in day 2 EpiLCs (Fig. 2e) and mESCs (Supplementary Fig. 2e). This was also shown upon STM2457 treatment in EpiLCs (Supplementary Fig. 2f). Moreover, depletion of m^6^A increased *Eras* mRNA transcript stability (Fig. 2f), in line with elevated *Eras* expression (Fig. 2c). *Oct4* or *GFP* transcripts which are not subject to m^6^A modification^24^ (Supplementary Fig. 2g), displayed no m^6^A depletion dependent transcript stability effects (Supplementary Fig. 2h, i). Hence, *Eras* transcripts are m^6^A decorated and can undergo m^6^A depletion. Consequently, the increased transcript stability elevates *Eras* levels.

To characterize if *Eras* transcript clearance is regulated by m^6^A reader redundancy or specificity mechanisms^23^, we implemented *Ythdf1, 2* or *3* siRNA knock-down assays during EpiLC induction. *Eras* mRNA level was increased upon siRNA mediated knock-down of *Ythdf1* or *2*, but not *Ythdf3* (Fig. 2g). *Ythdf1* and *2* depletion showed additive increases in mRNA expression (Fig. 2g) and transcript stability (Supplementary Fig. 2j). Correspondingly, ERAS and pAKT protein levels were highest upon siRNA mediated knock-down of *Ythdf1* and *2* (Supplementary Fig. 2k), but not *Ythdf3* (Supplementary Fig. 2l). Hence, *Ythdf1 and 2* coordinate m^6^A dependent *Eras* regulation in EpiLCs.

To confirm that elevated *Eras* levels suppress germline entry, we conducted a siRNA mediated *Eras* knock-down in m^6^A depleted day 2 EpiLCs (Supplementary Fig. 2m, n) and performed germline induction. *Eras* levels remained low in mPGCLCs (Supplementary Fig. 2o). *Eras* knock-down rescued germline entry efficiency in m^6^A depleted cells. This was reflected by an increased proportion of SSEA1 and CD61 double positive (Fig. 2h, Supplementary Fig. 2p) or *Blimp1*-GFP reporter positive (Supplementary Fig. 2q) day 4 mPGCLC spheres. RT-qPCR analysis further confirmed restored expression levels of key germline genes, including *Blimp1, Prdm14, Tfap2c, Nanos3, Stella* and *Dnd1* (Fig. 2i).

### Depletion of m^6^A and Eras Upregulation Impair Germline Entry by Reducing H3K27 Trimethylation

We found that ERAS protein expression upon m^6^A depletion was accompanied by greater phosphorylation of AKT on T308 and S473 residues (Fig 3a). *Eras* siRNA mediated knock-down in m^6^A depleted cells diminished pAKT activation (Fig. 3a). The increased proportion of *Oct4*ΔPE::GFP positive day 2 EpiLCs upon m^6^A depletion could be restored either by siRNA mediated knock-down of *Eras* (Fig. 3b, Supplementary Fig.3a), *Akt* knock-down (Supplementary Fig. 3b-d) or the pharmacological inhibition of PI3K (Fig. 3c, Supplementary Fig. 3e). In line with ERAS-pAKT dependent regulation, the AKT specific inhibitor MK2206 also reduced the proportion of *Oct4*ΔPE::GFP positive day 2 EpiLCs (Supplementary Fig. 3f, g) and rescued germline entry deficiency in m^6^A depleted cells (Supplementary Fig. 3h). Epigenetic remodelling during the establishment of formative pluripotency appears essential for germline competence^30,31,47-50^ but remains poorly defined. Increased pAKT was previously shown to negatively regulate EZH2, the enzymatically active component of the Polycomb Repressive Complex 2 (PRC2)^51,52^. Depletion of m^6^A reduced EZH2 mRNA and protein levels as well as global abundances of the H3K27me3 histone mark (Fig. 3d). In comparison, H3K9me2, H3K27ac and H3K4me1 histone marks associated with active or poised enhancers^30,47,53,54^ remained unchanged (Fig. 3d). Chromatin-immunoprecipitation-qPCR (ChIP-qPCR) confirmed reduced H3K27me3 deposition on the *Oct4* distal enhancer locus, but not H3K27ac, H3K4me1 or H3K9me2 levels (Fig. 3e).

**Figure 3.**
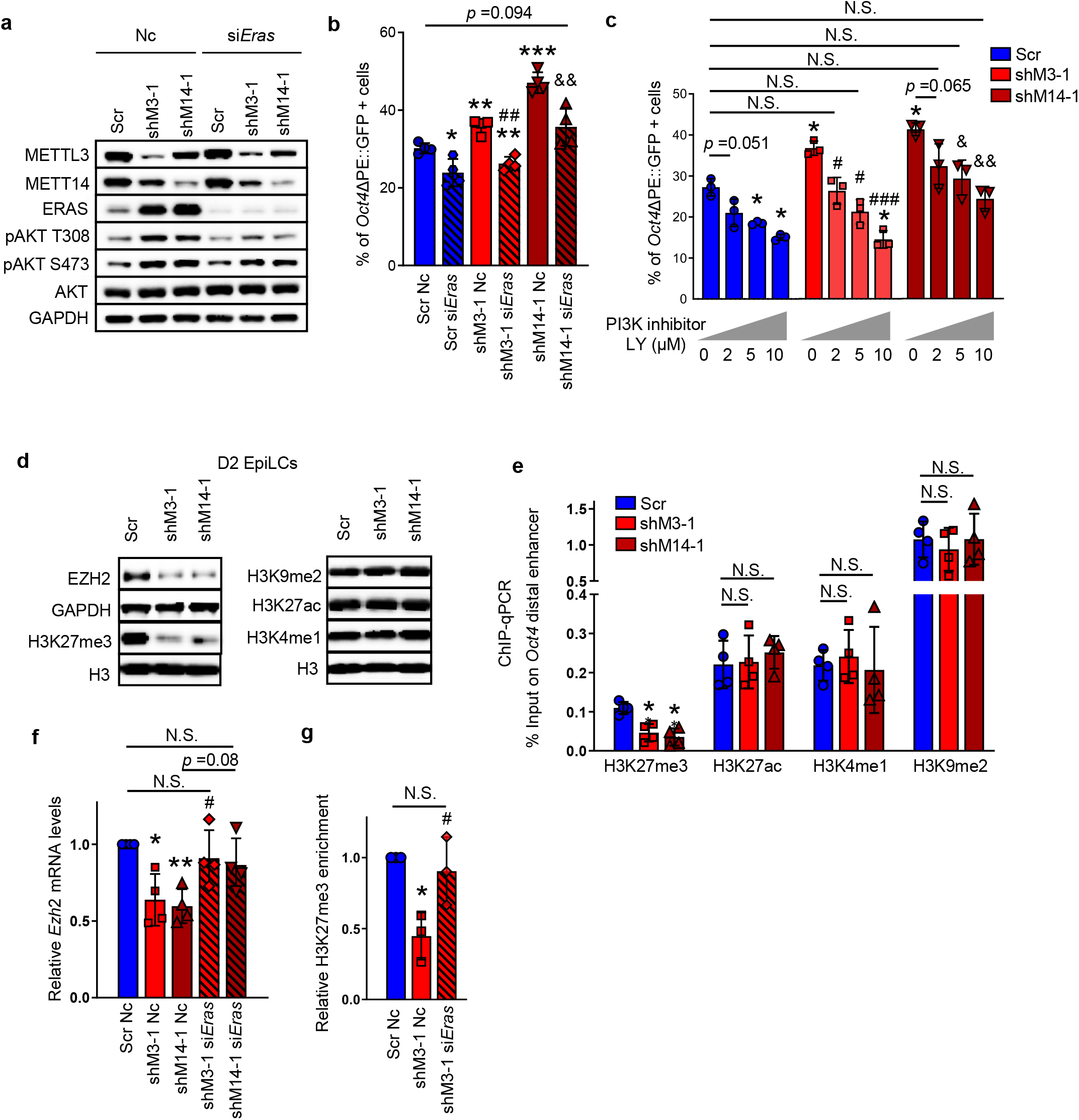
Depletion of m^6^A and *Eras* Upregulation Impair Germline Entry by Reducing H3K27 Trimethylation. **a-c**, Oct4ΔPE::GFP ESCs were transduced with scramble control (Scr), *Mettl3* (shM3-1) or *Mettl14* (shM14-1) shRNA before EpiLC induction. Day 0 EpiLCs were either (**a, b**) transfected with negative control (Nc) or *Eras* (si*Eras*) siRNA, or (**c**) treated with DMSO or PI3K inhibitor LY294002 (LY). (**a**) Immunoblotting analysis and (**b, c**) graphs summarizing FACS analyses of *Oct4*ΔPE::GFP positive cells in day 2 EpiLCs. Mean ± SD. Two-tailed, paired t-test: vs Scr or Scr Nc, \*\*\**p* <0.001; \*\**p* <0.01; \**p* <0.05; vs shM3-1 or shM3-1 Nc, ^###^*p* <0.001; ^##^*p* <0.01; ^#^*p* <0.05; vs shM14-1 or shM14-1 Nc, ^&&^*p* <0.01; ^&^*p* <0.05; N.S., Non-significant. See Figure S2m for experimental protocol and S3a and Supplementary Files for representative FACS plots. **d-e**, *Oct4*ΔPE::GFP ESCs were transduced with scramble control (Scr), *Mettl3* (shM3-1) or *Mettl14* (shM14-1) shRNA before EpiLC induction. (**d**) Immunoblotting and (**e**) ChIP-qPCR at the *Oct4* distal enhancer locus in day 2 EpiLCs. Mean ± SD. N =4; two-tailed, paired t-test: vs Scr, * *p*<0.05; N.S., Non-significant. **f-g**, ESCs were transduced with scramble control (Scr), *Mettl3* (shM3-1) or *Mettl14* (shM14-1) shRNA before EpiLC induction. Day 0 EpiLCs were transfected with negative control or *Eras* (si*Eras*) siRNA. (**f**) qPCR in *Blimp1*-GFP day 2 EpiLCs (N =4). (**g**) ChIP-qPCR at the *Oct4* distal enhancer locus in *Oct4*ΔPE::GFP day 2 EpiLCs (N =3). Mean ± SD. Two-tailed, paired t-test: vs Scr Nc, \*\**p* <0.01; \**p* <0.05; vs shM3-1 Nc, ^#^*p* <0.05; N.S., Non-significant See also Figure S2m for experimental protocol.

We sought to define whether ERAS was responsible for H3K27me3 level modulation upon m^6^A depletion. *Eras* siRNA mediated knock-down in m^6^A depleted cells restored *Ezh2* expression (Fig. 3f), re-instated global H3K27me3 levels (Supplementary Fig. 3i) and H3K27me3 marks on the *Oct4* distal enhancer (Fig. 3g). Next, we aimed to determine if ERAS upregulation was able to phenocopy these epigenetic changes in the absence of m^6^A depletion. However, while ERAS over-expression in wild-type cells activated pAKT, no change in H3K27me3 deposition was found (Supplementary Fig. 3j). Neither the increased proportion of *Oct4*ΔPE::GFP positive day 2 EpiLCs (Supplementary Fig. 3k) nor germline entry deficiency associated with m^6^A depletion (Supplementary Fig. 3l) could be phenocopied by ERAS over-expression in wild-type cells. This suggested that m^6^A depletion and/or ERAS found in an m^6^A depletion setting, modulates the transcription of *Ezh2* and consequently, reduces H3K27me3 abundances. Intriguingly, co-immunoprecipitation analyses in wild-type and m^6^A depleted EpiLCs overexpressing *Eras*, show that ERAS and EZH2 proteins can physically interact only upon m^6^A depletion (Supplementary Fig. 3m). Hence, decreasing m^6^A abundances establishes unique molecular conditions whereby m^6^A depletion and *Eras* activation may repress *Ezh2* transcription and potentially, ERAS may also physically modulate EZH2 activity in EpiLCs.

### Depletion of m^6^A and Reduced H3K27 Trimethylation Deregulates EpiLC Gene Expression

We showed earlier that depletion of m^6^A removed H3K27 trimethylation at the *Oct4* distal enhancer. Therefore, we sought to characterize the genome-wide changes in H3K27me3 localization by applying ChIP-seq to profile the H3K27me3 landscape in control and m^6^A depleted (STM2457 treated) day 2 EpiLCs (Supplementary Fig. 4a). In addition, we performed RNA-seq to monitor the associated changes in gene expression and cell state regulation (Supplementary Fig. 4b). Previous work established that in some cell types, the majority of H3K27me3 exists as dispersed, low abundance signal outside of high enrichment areas^55^. We identified a profound and general redistribution of H3K27me3 whereby m^6^A depletion resulted in a global reduction of dispersed, low abundance H3K27 trimethylation existing in large parts of the genome (Fig. 4a). As previously indicated, this also included the *Oct4* distal and proximal enhancers (Fig. 4b). ChIP-qPCR further affirmed our ChIP-seq profiling of *Oct4* regulatory regions (Fig. 4c).

**Figure 4.**
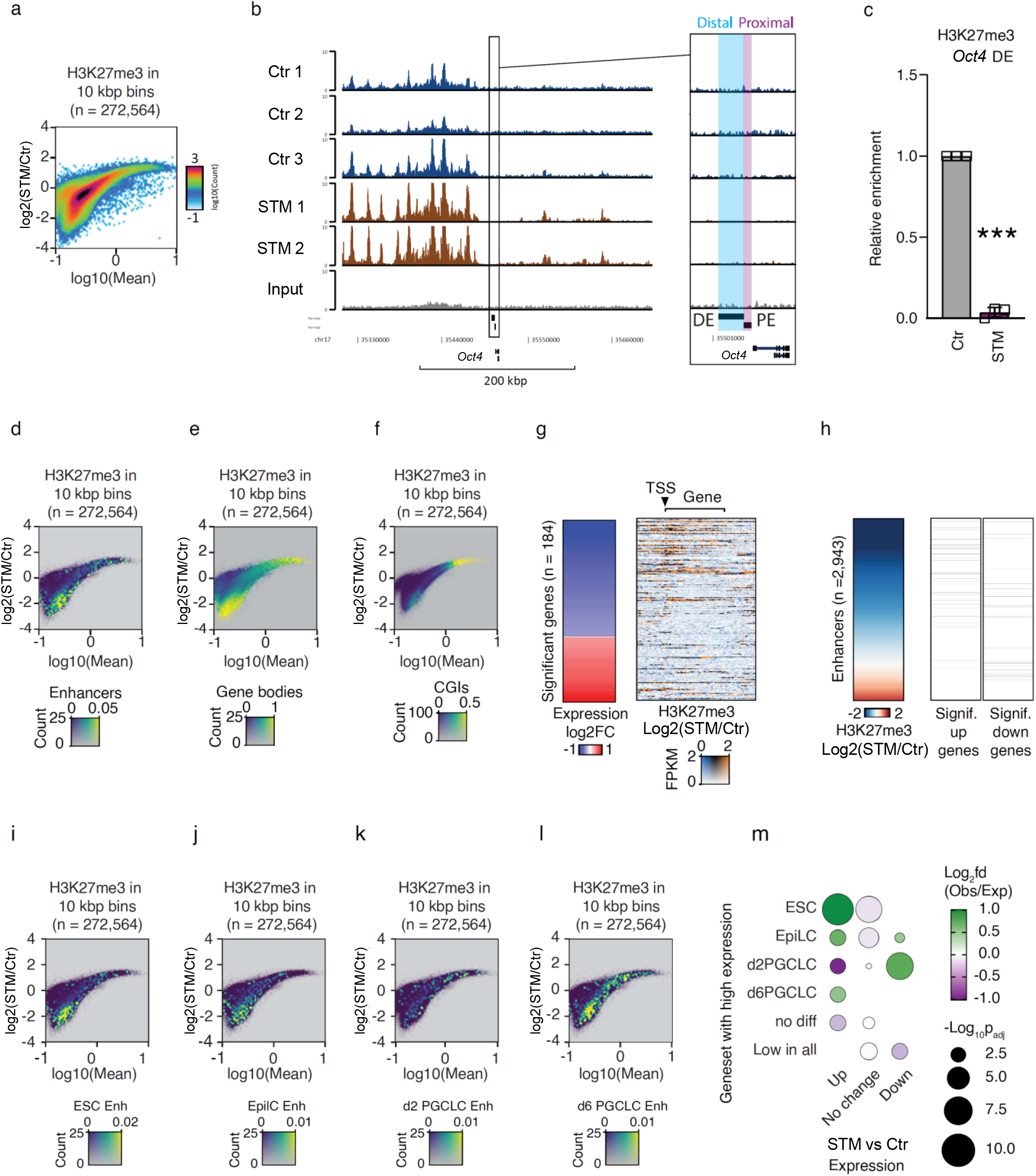
Depletion of m^6^A and Reduced H3K27 Trimethylation Deregulates EpiLC Gene Expression. **a-m**, *Blimp1*-GFP ESCs were induced into EpiLCs, and day 0 EpiLCs were treated with DMSO (Ctr) or METTL3 inhibitor STM2457 (STM) for 2 days, followed by ChIP sequencing (**a-b, d-l**), qPCR (**c**) and RNA sequencing (**g-h, m**) analyses. **a**, MA plot showing the relationship between mean H3K27me3 signal (X-axis) and log_2_ fold difference in H3K27me3 signal (Y-axis). The H3K27me3 signal was analyzed in 10 kbp bins throughout the genome, values were normalized to Fragments per Kilobasepair per Million reads (FPKM), and averaged across biological replicates. **b**, Genome-browser tracks showing H3K27me3 peaks. Values are FPKM normalized, and the insert shows a zoom in on the genomic area containing the *Oct4* distal enhancer (DE; blue) and proximal enhancer (PE; purple). **c**, ChIP-qPCR showing H3K27me3 signal changes in *Oct4* distal enhancer. N =3; two-tailed, paired t-test: \*\*\**p* <0.001. **d-f**, Coloured MA-plots showing the three-way relationship between the frequency of the indicated genomic features (colour-coded) as well as mean H3K27me3 signal (X-axis) and log_2_ fold difference in H3K27me3 signal (Y-axis). The H3K27me3 signal was analyzed in 10 kbp bins throughout the genome, values were normalized to FPKM, and averaged across biological replicates. (**d**), Putative enhancers; (**e**), Gene bodies; (**f**), CpG islands (CGIs). Enhancer coordinates were obtained from the Kurimoto identified enhancers in ESCs, EpiLCs, and day 2/6 PGCLCs ^29^. **g**, Heatmaps showing the log_2_ fold difference in expression (left) for genes significantly affected upon STM2457 treatment as well as the log_2_ fold differences in H3K27me3 levels at the gene body and surrounding loci (right). Changes in FPKM-normalized H3K27me3 levels are colored blue for reduced, black for unchanged, and orange for increased. Opacity is controlled by mean H3K27me3 levels in the two conditions. See Supplementary Table 5 for the gene expression changes and Figure S4 c-h for individual H3K27me3 heatmaps. TSS, Transcription start site. **h**, Heatmaps showing log_2_ fold differences in FPKM-normalized H3K27me3 levels at 2,943 enhancers identified according to cell states (ESC, EpiLC, and day 2/6 PGCLC) ^29^ (left) as well as the overlap of significantly upregulated (middle) or downregulated (right) genes nearest to each putative enhancers. See Figure S4 l-q for individual H3K27me3 heatmaps in these enhancers. **i-l**, Coloured MA-plots showing the three-way relationship between the frequency of the indicated genomic features (color-coded) as well as mean H3K27me3 signal (X-axis) and log_2_ fold difference in H3K27me3 signal (Y-axis). The H3K27me3 signal was analyzed in 10 kbp bins throughout the genome, values were FPKM normalized, and averaged across biological replicates. H3K27me3 changes in stage-specific active enhancer subgroups (Kurimoto classification^29^) for: ESC (**i**), EpiLC (**j**), d2 PGCLC (**k**) and D6 PGCLC (**l**). **m**, Bubble plots showing the relative abundance and significance compared to random expectance between groups of genes that are significantly affected or unaffected (X-axis) and previously identified as highly expressed in indicated subtypes of cells^29^ (Y-axis). Bubble colour reflects the log_2_ fold difference between the observed overlap compared to the expected if all categories are equally likely, whereas bubble sizes are proportional to the −log_10_ significance of the overlap. P-values are Benjamini-Hochberg adjusted for multiple testing. See Supplementary Table 6 for detailed gene list.

Next, we examined the frequency of reported putative enhancers^29^, as well as global CpG-islands and gene bodies in relation to changes in H3K27me3. This revealed a general depletion of dispersed, low abundance H3K27me3 signal at enhancers (Fig. 4d), wide-spread H3K27me3 depletion across gene-bodies (Fig. 4e) and gain of H3K27me3 at regions coinciding with CpG islands (Fig. 4f). Given the compositional nature of ChIP-seq data^56^, a massive reduction in dispersed, low abundance signal in gene bodies led to an apparent increase in reads sequenced from the remaining high abundance H3K27 trimethylation^55^. To explore transcriptional changes associated with alterations in H3K27me3, we identified significantly differentially expressed genes and ordered these according to fold change in expression. Remarkably, reduced H3K27 trimethylation upon m^6^A depletion, including at low abundance regions, correlated with significantly increased gene expression (Fig. 4g and Supplementary Fig. 4b-j). Such an inverse relationship is consistent with the repressive role of H3K27me3^54,55^ and reinforces the functional importance of dispersed, low abundance H3K27 trimethylation regions in gene regulation.

This association also persisted when examining global enhancer H3K27me3 levels and the expression changes of their nearest genes. Enhancers with reduced H3K27 trimethylation upon m^6^A depletion generally existed near genes with significantly increased expression (Fig. 4h and Supplementary Fig. 4k-s). Next, we examined sets of previously reported putative enhancers partitioned into subsets of different embryonic cell states by Kurimoto et al., (2021)^29^. This included the mESC, EpiLC as well as day 2 and day 6 PGCLC states. Nearly all mESC (Fig. 4i) and EpiLC (Fig. 4j) specific gene enhancers showed prominent H3K27me3 hypomethylation. A modest trend of H3K27me3 hypermethylation was found in day 2 PGCLC enhancers (Fig. 4k). The day 6 PGCLC cohort showed a large bimodal deregulation pattern, with prominent hypo- and hypermethylation of H3K27me3 (Fig. 4l). We examined these H3K27me3 patterns in relation to gene expression characterizing mESC, EpiLC as well as day 2 and day 6 PGCLC cell states^29^. Upregulation of m^6^A-depleted genes was significantly enriched among those previously categorized as mESC and EpiLC genes (Fig. 4m). Conversely, repressed genes were significantly enriched among those previously categorized as day 2 PGCLC genes (Fig. 4m). Together, our data indicates that m^6^A depletion promotes a reduction of global, low abundance H3K27 trimethylation in EpiLCs, with profound changes also on enhancers. We propose that these alterations facilitate unscheduled naïve pluripotent and epiblast-like gene upregulation, interfering with germline program establishment. Other m^6^A dependent mechanisms, including changes to transcript stability or translation, may also contribute to the observed alterations in gene expression.

### Depletion of m^6^A Promotes Eras Dependent Germline Entry Deficiency in Mouse FS cells

Recent work has identified formative state self-renewal conditions that rely on endogenous FGF signaling^2^. Initially, *Mettl3* or *Mettl14* knock-down was performed in 2i+LIF mESCs, followed by FS cell induction (Supplementary Fig. 5a). Here, *Eras* upregulation (Supplementary Fig. 5b) was accompanied by an increase in the proportion of *Oct4*ΔPE::GFP positive cells at day 2 of FS cell state entry (Fig. 5a, b). However, this effect ceased by day 4 (Fig. 5a), indicating that m^6^A depletion does not suppress but delays enhancer silencing kinetics during formative cell state acquisition. In line with this, ATAC-seq^2^ shows that FS cells eventually adopt proximal enhancer usage to drive *Oct4* expression (Supplementary Fig. 5c).

**Figure 5.**
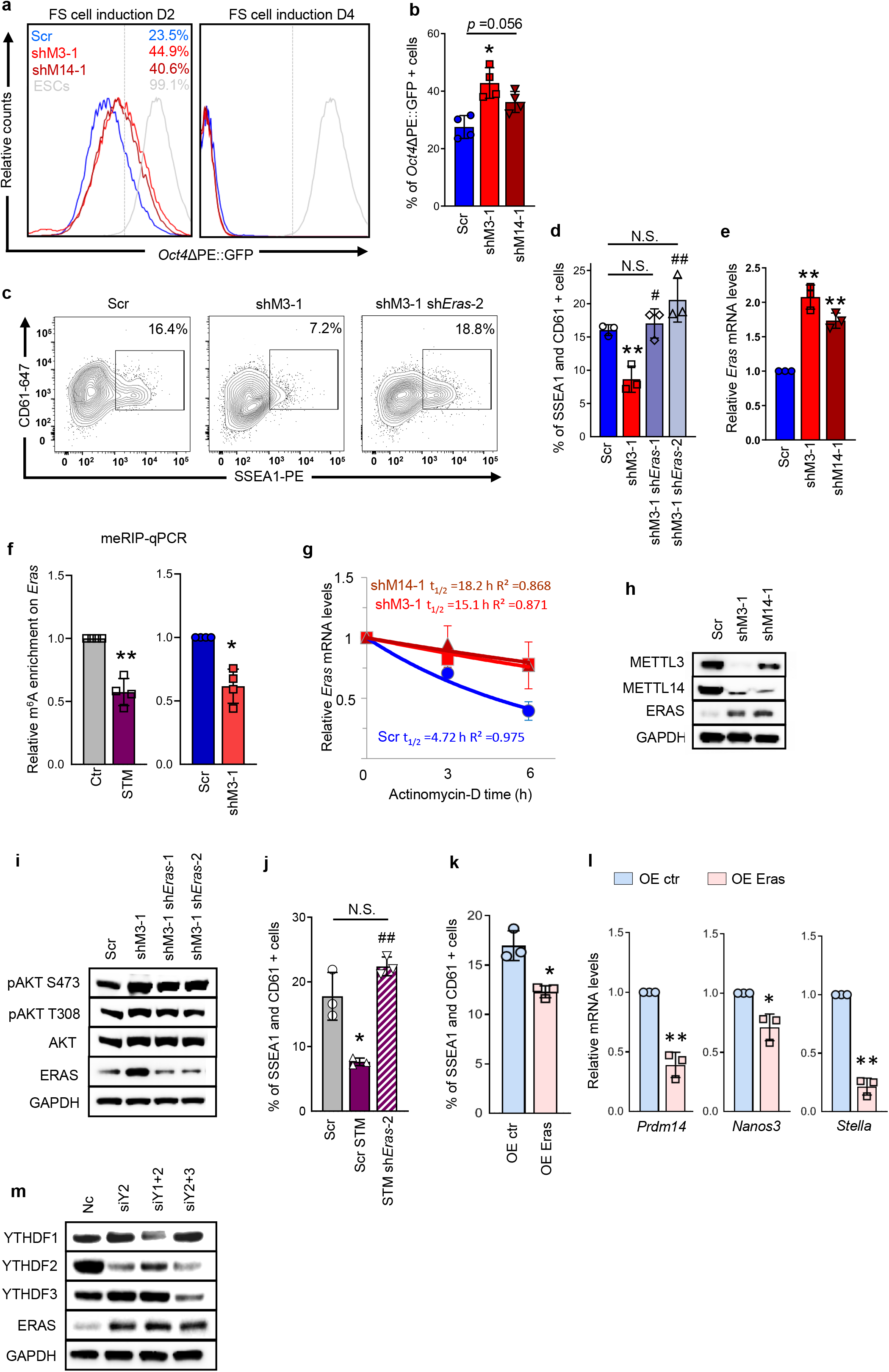
Depletion of m^6^A Promotes *Eras D*ependent Germline Entry Deficiency in Mouse FS cells. **a**, FACS analysis showing the percentage of *Oct4*ΔPE::GFP positive cells at day 2 and day 4 of Formative Stem (FS) cell induction from ESCs transduced either with scramble control (Scr), *Mettl3* (shM3-1) or *Mettl14* (shM14-1) shRNA. Dashed line separates positive gating. Solid gray line represents the GFP positive control. See Figure S4a for experimental protocol. **b**, Graph summarizing day 2 FACS data in (**a**). Mean ± SD. N =4; two-tailed, paired t-test: vs Scr, \**p* <0.05. **c-j**, *Blimp1-*GFP reporter FS cells were treated with DMSO or METTL3 inhibitor STM2457, or transduced with scramble control, *Mettl3* (shM3-1), *Mettl14* (shM14-1) or *Mettl3* and *Eras* (shM3-1 sh*Eras*-1 or shM3-1 sh*Eras*-2) shRNA before PGCLC specification. Mean ± SD. Two-tailed, paired t-test: vs Scr or Ctr, \*\**p* <0.01; vs shM3-1 or Scr STM, ^##^*p* <0.01; ^#^*p* <0.05; N.S., Non-significant. See Figure S5d for the experimental protocol. **c**, FACS analysis showing the percentage of SSEA1 and CD61 double positive cells in day 4 PGCLC spheres. **d**, Graph summarizing effects in (**c**). N =3. *e, Eras* qPCR analysis in FS cells. N =3. **f**, meRIP-qPCR showing m^6^A enrichment on *Eras* mRNA. N =4. **g**, qPCR showing *Eras* mRNA half-life (t_1/2_) in FS cells. See Supplementary Table 2 for statistical details. **h**, Immunoblots showing METTL3, METTL14 and ERAS levels in FS cells. **i**, Immunoblots showing ERAS, AKT and pAKT levels in FS cells. **j**, Graph summarizing FACS data showing the percentage of SSEA1 and CD61 double positive cells in PGCLC. N =3. See Supplementary Files for representative FACS plots. **k-l**, R1 FS cells transduced with either control (OE Ctr) or ERAS-overexpressing (OE Eras) lentivirus were induced into PGCLCs for 4 days. **k**, Graph summarizing FACS data showing the percentage of SSEA1 and CD61 double positive cells in PGCLC spheres. See Supplementary Files for representative FACS plots. **l**, qPCR showing germline marker gene expression in PGCLC spheres. **m**, Immunoblots from *Blimp1*-GFP FS cells transfected with negative control (Nc), *Ythdf2* (siY2), *Ythdf1* and *Ythdf2* (siY1+2) or *Ythdf2* and *Ythdf3* (siY2+3) siRNA.

Depletion of m^6^A in expanded, self-renewing FS cells (Supplementary Fig. 5d, e) showed no visible effects on cell growth or morphology (Supplementary Fig. 5f). However, we found significantly impaired germline entry efficiency (Fig 5c, d, Supplementary Fig. 5g). As in EpiLCs, an increase in *Eras* expression was observed in m^6^A depleted FS cells (Fig. 5e). *Fgf4* or *Fgf5* upregulation, previously reported upon m^6^A depletion in metastable pluripotent Serum/LIF mESCs^57^, remained unchanged (Supplementary Fig. 5h). We confirmed by meRIP-qPCR, a significant m^6^A depletion on the 3’ end of *Eras* mRNA (Fig. 5f). This corresponded with increased *Eras* transcript stability (Fig. 5g). Changes to *Eras* mRNA expression were consistent with elevated ERAS protein levels (Fig. 5h) as well as AKT T308 and S473 phosphorylation (Fig. 5i). Germline entry efficiency could be restored by shRNA mediated knock-down of *Eras* (Fig. 5c, d, i, Supplementary Fig. 5g) or modulation of *Eras* dependent signaling in m^6^A depleted cells (Supplementary Fig. 5i). These effects were recapitulated by METTL3 inhibition using STM2457 treatment (Fig. 5j, Supplementary Fig. 5j-l). Short-term STM2457 withdrawal (Supplementary Fig. 5m) also resulted in *Eras* modulation (Supplementary Fig. 5n) and restored germline entry efficiency (Supplementary Fig. 5o). Furthermore, *Eras* over-expression in wild-type FS cells significantly impaired germline entry (Fig. 5k), albeit at supraphysiological levels (Supplementary Fig. 5p). This was accompanied by a significant depletion in the expression of key germline genes (Fig. 5l). The inability to reproduce this effect in wild-type EpiLCs (Supplementary Fig. 3j-l) may reflect differences in formative cell states or technical challenges in characterizing a transient EpiLC system. Finally, while *Eras* transcript clearance is synergistically regulated by YTHDF1 and YTHDF2 in EpiLCs, *Eras* mRNA stability and pAKT activation in FS cells is modulated by YTHDF2 alone (Fig. 5m, Supplementary Fig. 5q, r). Preference for different YTHDFs to modulate *Eras* levels in distinct formative cell populations, suggests a degree of mechanistic plasticity in m^6^A dependent gene regulation.

### Depletion of m^6^A Promotes Mesodermal Differentiation in Mouse FS Cells

Deregulated gene expression during EpiLC state acquisition likely occurs due to both epigenetic and transcript stability dependent changes in m^6^A depleted cells. In expanded, self-renewing FS cells, decreased m^6^A abundances by shRNA knock-down of *Mettl3* or STM2457 mediated METTL3 inhibition, induced a cell population with unscheduled upregulation of *Blimp1*. To characterize this effect, we employed flow cytometry using a *Blimp1*-GFP reporter cell line (Fig. 6a, b, Supplementary Fig. 6a, b) and protein level analyses (Supplementary Fig. 6c, d). The accumulation of Blimp1-GFP positive cells was reversible by STM2457 withdrawal (Supplementary Fig. 6b). shRNA mediated knock-down of *Eras* (Fig. 6a, b, Supplementary Fig. 6c-e) or its downstream signaling inhibition (Supplementary Fig. 6f, g) significantly reduced, but did not eliminate the *Blimp1*-GFP positive cell fraction. Persistence of these cells in culture was likely due to reduced m^6^A deposition found on the 3’ untranslated region (UTR) of *Blimp1* mRNA upon m^6^A depletion (Fig. 6c, Supplementary Fig. 6h). This was associated with elevated *Blimp1* transcript stability (Fig. 6d).

**Figure 6.**
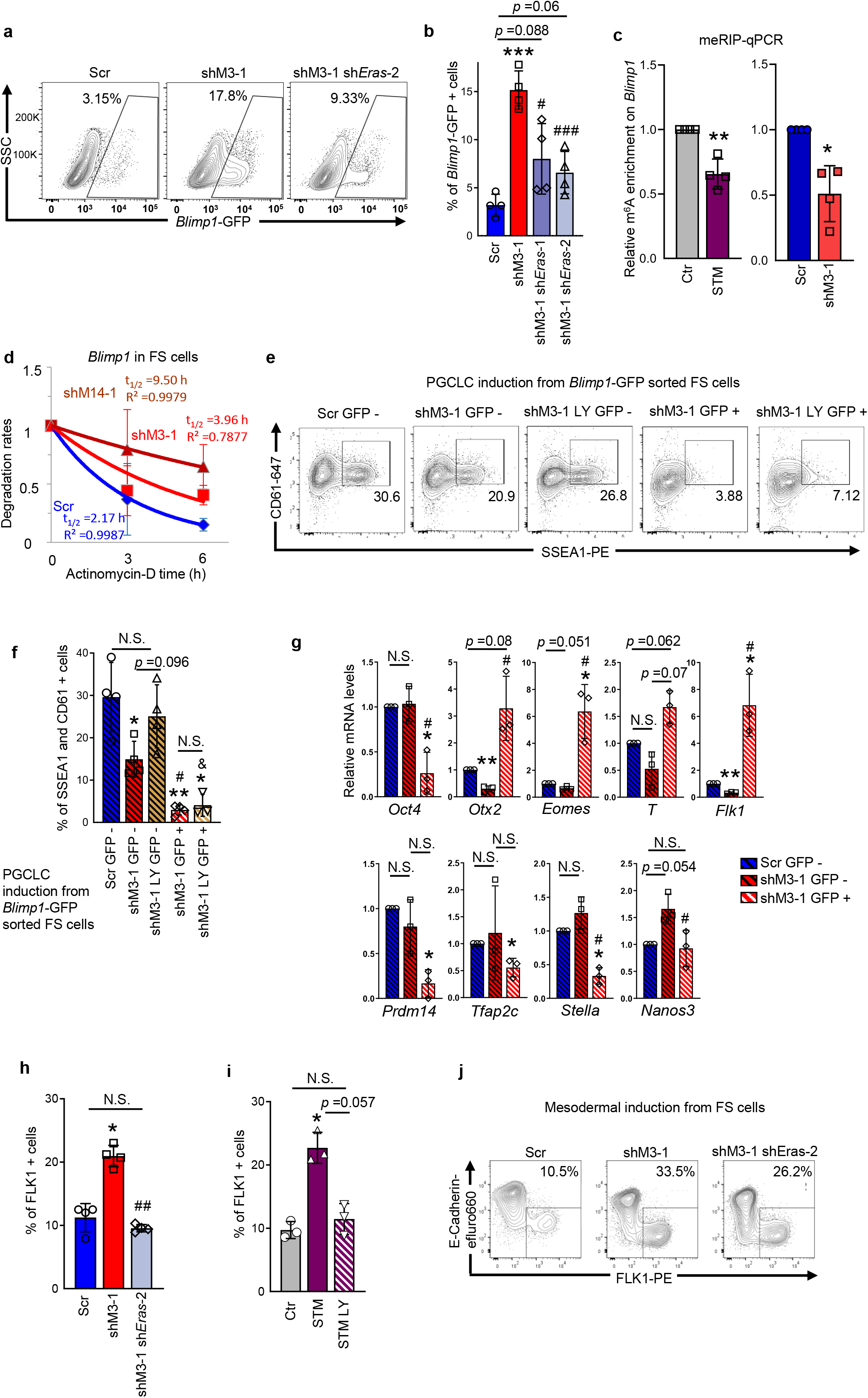
Depletion of m^6^A Promotes Mesodermal Differentiation in Mouse FS Cells. **a-d**, *Blimp1*-GFP reporter FS cells were either treated with DMSO or METTL3 inhibitor STM2457 for 2 days, or transduced with scramble control (Scr), *Mettl3* (shM3-1), *Mettl14* (shM14-1) or *Mettl3* and *Eras* (shM3-1 sh*Eras*-1 or shM3-1 sh*Eras*-2) shRNA for 10 days. Mean ± SD. Two-tailed, paired t-test: vs Scr or Ctr, \**p* <0.05; \*\**p* <0.01; \*\*\**p* <0.001; vs shM3-1, ^###^*p* <0.001; ^#^*p* <0.05. **a**, FACS analysis showing the percentages of *Blimp1*-GFP positive FS cells. SSC, side scatter parameter. **b**, Graph summarizing effects in (**a**). N =4. **c**, meRIP-qPCR showing m^6^A enrichment on *Blimp1* mRNA. N =4. **d**, qPCR showing *Blimp1* mRNA half-life (t_1/2_). N =3. See Supplementary Table 2 for statistical details. **e-g**, *Blimp1*-GFP reporter FS cells were transduced with scramble control (Scr) or *Mettl3* (shM3-1) shRNA, then either (**g**) sorted into GFP positive and negative cell fractions, or (**e-f**) treated with DMSO or PI3K inhibitor LY294002 (LY) followed by GFP positive and negative cell sorting for PGCLC induction. Mean ± SD. N ≥3; two-tailed, paired t-test: vs Scr GFP−, \*\**p* <0.01; \**p* <0.05; vs shM3-1 GFP−, ^#^*p* <0.05; vs shM3-1 LY GFP−, ^&^*p* <0.05; N.S., Non-significant. **e**, FACS plots showing the percentages of SSEA1 and CD61 double positive cells in PGCLC spheres. **f**, Graph summarizing effects in (**e**). **g**, qPCR showing gene expression changes in sorted FS cell fractions. **h-i**, Graph summarizing FACS data showing the percentage of FLK1 positive cells in day 4 PGCLC spheres from *Blimp1*-GFP FS cells with different treatments. (**h**) FS cells were transduced with scramble control (Scr), *Mettl3* (shM3-1) or *Mettl3* and *Eras* (shM3-1 sh*Eras*-2) shRNA; (**i**) FS cells were treated with DMSO (Ctr), METTL3 inhibitor STM2457 (STM) or STM2457 and LY294002 (STM LY). Mean ± SD. N =4; two-tailed, paired t-test: vs Scr or Ctr, \**p* <0.05; vs shM3-1, ^##^*p* <0.01; N.S., Non-significant. See Supplementary Files for representative FACS plots. **j**, *Blimp1*-GFP reporter FS cells transduced with scramble control (Scr), *Mettl3* (shM3-1) or *Mettl3* and *Eras* (shM3-1 sh*Eras*-2) shRNA, were differentiated into the mesodermal lineage. FACS analysis showing the percentage of FLK1 positive and E-Cadherin negative mesodermal cells. See Figure S6r for experimental repeats.

To assess the germline competence of emergent cell sub-populations, fluorescence sorted m^6^A depleted *Blimp1*-GFP positive and negative cells were compared against the control *Blimp1*-GFP negative cell fraction (Supplementary Fig. 6i-j). Control *Blimp1*-GFP negative cells retained robust competence for germline entry. m^6^A depleted, *Blimp1*-GFP negative cells showed germline entry deficiency, rescuable by the inhibition of *Eras* dependent PI3K signalling (Fig. 6 e, f). By comparison, *Blimp1*-GFP positive, m^6^A depleted cells lacked germline competence and could not be rescued by signaling modulation (Fig. 6e, f). Therefore, we sought to determine why in m^6^A depleted FS cells, BLIMP1, a TF required for mouse germ cell regulatory network establishment^5-7^, was associated with germline entry deficiency. Gene expression analysis showed that sorted *Blimp1*-GFP positive FS cells displayed high *Otx2* co-expression and downregulation of *Oct4* (Fig. 6g). Mesendodermal markers like *Eomes* (Fig 6g, Supplementary Fig. 6k), *T* (Fig. 6g), and *Flk1* (Fig 6g, Supplementary Fig. 6l-n) became upregulated, while germline markers *Prdm14, Tfap2c, Stella* and *Nanos3* remained repressed (Fig. 6g). These results indicated that depletion of m^6^A promoted spontaneous differentiation partiality toward mesendoderm at the expense of germline competence in FS cells.

In line with this, PGCLC spheres established from m^6^A depleted FS cells showed increased expression of mesodermal marker *Flk1* (Supplementary Fig. 6o) and a higher proportion of FLK1 surface marker positive cells (Fig. 6h, i). The FLK1 positive fraction could be depleted by *Eras* knock-down (Fig. 6h) or inhibition of *Eras* signalling (Fig 6i, Supplementary Fig. 6p). Upregulated protein levels of mesodermal genes MESP1 and MIXl1 were also depleted by *Eras* knock-down (Supplementary Fig. 6q). In contrast, endodermal markers FOXA2 and GATA4 remained unchanged (Supplementary Fig. 6q). Further, directed differentiation of m^6^A depleted FS cells showed over two-fold increase in mesodermal lineage commitment (Fig. 6j, Supplementary Fig. 6r). *Eras* knock-down significantly reduced this differentiation partiality (Fig. 6j, Supplementary Fig. 6r), suggesting it can facilitate early mesodermal lineage specification in the epiblast where its expression is mostly confined^46^.

### Depletion of m^6^A Impedes Human Germline Entry and Promotes Neuroectoderm

*ERAS* is not expressed in human embryonic development^58^. To define the effects of decreasing m^6^A abundances on human germline competence, we examined hPSCs adapted to 4i conditions^14^. Consistently with mouse data, depletion of m^6^A by STM2457 treatment impaired germline entry. This was quantified by the proportion of EpCAM-APC, CD49f-BV421 fluorescent surface marker double positive cells at day 4 of hPGCLC induction (Fig. 7a, b). Upon m^6^A depletion, human 4i cells showed upregulation of *OTX2* levels and an increase in the proportion of *OTX2*-YFP positive reporter cells (Fig. 7c-e, Supplementary Fig. 7a). From the gene panel examined, germline markers *STELLA* and *DDX4* levels were also elevated (Fig. 7c, Supplementary Fig. 7b), consistently with their increased transcript stabilities (Supplementary Fig. 7c). However, transcript stability did not account for OTX2 upregulation in the human 4i hPSC system (Supplementary Fig. 7c). Instead, depletion of m^6^A resulted in the phosphorylation of FGFR1 and stimulation of downstream pERK and pAKT signaling pathways in 4i hPSCs, but not in mouse support feeder cells (Fig. 7f, Supplementary Fig. 7d). Pharmacological inhibition of FGFR1 or MEK significantly depleted OTX2 levels (Supplementary Fig. 7d, e) while PI3K inhibition showed minor effects (Supplementary Fig. 7e, f). RT-qPCR analysis of FGF signaling ligands identified upregulation of *FGF5* and *FGF10* (Supplementary Fig. 7g), together with their increased transcript stabilities (Supplementary Fig. 7c). These findings highlight that in human formative pluripotency, FGFR1-pERK activation promotes OTX2 expression in response to m^6^A depletion.

**Figure 7.**
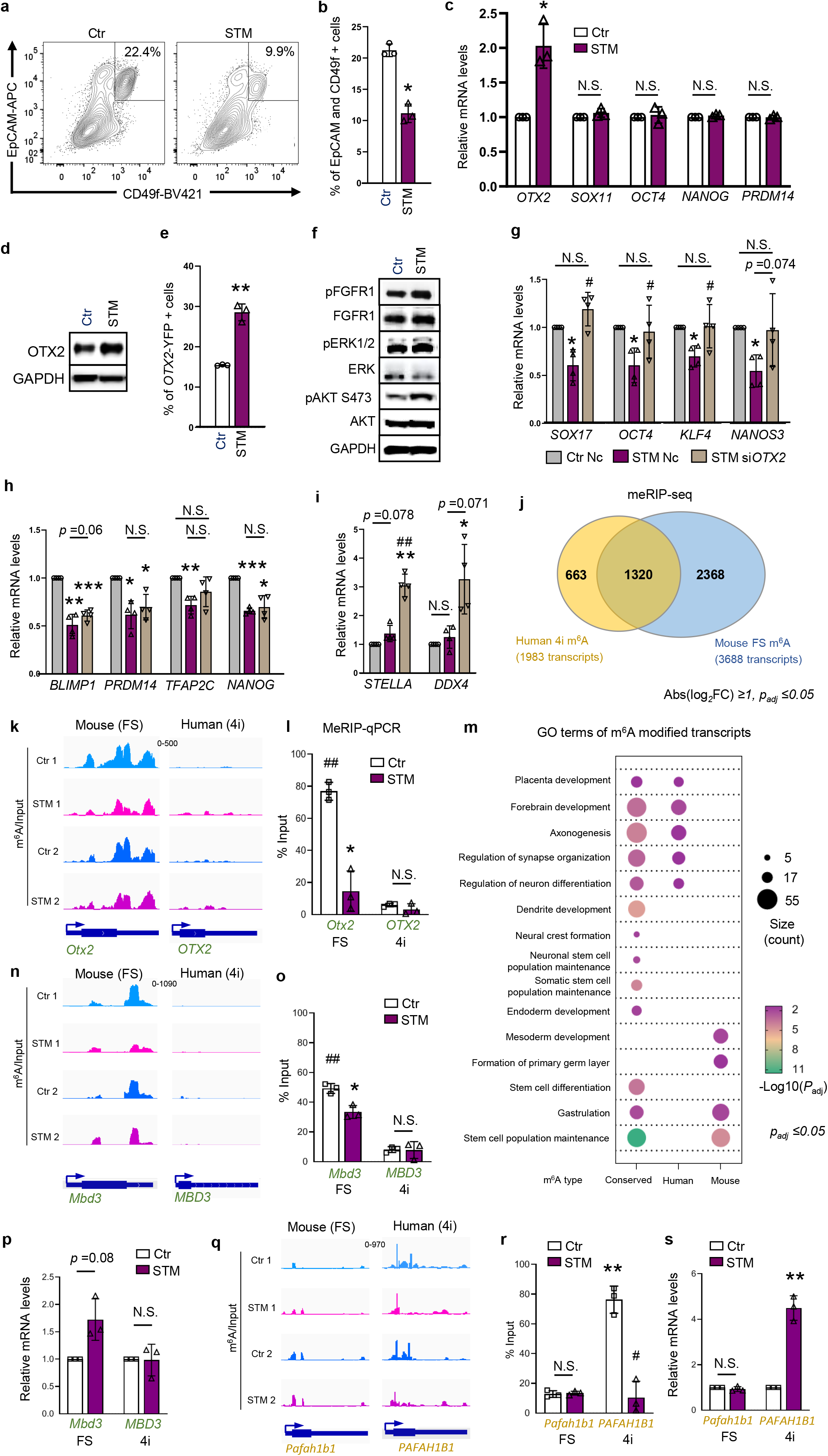
Depletion of m^6^A Impedes Human Germline Entry and Promotes Neuroectoderm. **a-f**, H9 wild-type or *OTX2*-YFP reporter 4i hPSCs were treated with DMSO (Ctr) or METTL3 inhibitor STM2457 (STM) for 3 days before hPGCLC induction. Mean ± SD. N =3; two-tailed, paired t-test: vs Ctr, \**p*<0.05; \*\**p* <0.01. **a**, Representative FACS analysis showing the percentage of human germline marker EpCAM and CD49f double positive cells in *OTX2*-YFP day 4 PGCLC spheres. **b**, Graph summarizing effects in (**a**). **c**, qPCR in H9 4i hPSCs showing expression changes of formative pluripotency markers. **d**, Immunoblots showing OTX2 expression in H9 4i hPSCs. **e**, Graph summarizing FACS analyses showing the percentages of *OTX2*-YFP positive cells in 4i hPSCs. See Figure S7a for representative FACS plots. **f**, Immunoblots showing changes in the phosphorylation of FGFR1, ERK and AKT in H9 4i hPSCs. **g-i**, qPCR showing mRNA changes for germline genes in day 4 PGCLC spheres, derived from H9 4i hPSCs treated with DMSO and negative control siRNA (Ctr Nc), STM2457 and negative control siRNA (STM Nc) or STM2457 and *OTX2* siRNA (STM si*OTX2*) for 3 days. (**g**) *OTX2* knock-down restored germline gene expression depleted by STM2457 treatment. (**h**) *OTX2* knock-down failed to restore germline gene expression. (**i**) *OTX2* knock-down promoted germline gene expression. Mean ± SD. N =4; two-tailed, paired t-test: vs Ctr, \*\*\**p* <0.001; \*\**p* <0.01; \**p* <0.05; vs STM, ^##^*p* <0.01;^#^*p* <0.05; N.S., Non-significant. **j-s**, Comparative m^6^A profiling in mouse and human formative stem cells. meRIP-seq and qPCR analyses in mouse R1 FS cells and human H9 4i hPSCs with DMSO (control) or STM2457 (STM) treatment. **j**, Venn diagram of m^6^A-modified transcripts shared between mouse and human formative stem cells. **k, n, q**, Representative genome browser tracks showing m^6^A enrichment (m^6^A-IP subtract Input signals) for mouse and human specific m^6^A-modified genes. See Supplementary Files for individual IGV tracks showing Input and m^6^A-IP peak signals. **l, o, r**, meRIP-qPCR validation of m^6^A modification levels in *OTX2, MBD3* and *PAFAH1B1* RNAs. **m**, Gene Ontology (GO) analysis of shared and mouse- and human-specific m^6^A transcripts. **p, s**, qPCR showing the RNA level changes in *MBD3* and *PAFAH1B1*.

OTX2 was identified as a repressor of both mouse^27^ and human^28,59^ germline entry. Correspondingly, an increase in the proportion of *OTX2*-YFP high cells found in m^6^A depleted hPGCLC spheres was mutually exclusive with CD49f-BV421 germ cell surface marker expression (Supplementary Fig. 7h). However, *OTX2* siRNA-mediated knock-down in m^6^A depleted 4i cells prior to hPGCLC induction did not rescue germline entry deficiency (Supplementary Fig. 7i, j). RT-qPCR analysis in day 4 hPGCLC spheres confirmed that m^6^A depletion culminated in the downregulation of key germline markers (*SOX17, OCT4, KLF4* and *NANOS3*) (Fig. 7g). While *OTX2* knock-down restored the expression of these markers in m^6^A depleted cells (Fig. 7g), the levels other germline genes (*BLIMP1, PRDM14, TFAP2C* and *NANOG*) remained low (Fig. 7h). This may explain why *OTX2* knock-down cannot rescue the impairment of germline entry upon m^6^A depletion (Supplementary Fig. 7j). Furthermore, knock-down of *OTX2* in m^6^A depleted 4i hPSCs elevated the expression of germline markers *STELLA* and *DDX4* in hPGCLC spheres (Fig. 7i). Hence, *OTX2* appeared to counterbalance the effects of increased transcript stabilities observed for *STELLA* and *DDX4* upon m^6^A depletion (Supplementary Fig. 7c). Perhaps low OTX2 levels moderate some human germline genes, facilitating correct dosage for the acquisition of germline competence. Finally, directed differentiation assays toward mesoderm, endoderm and neuroectoderm showed that m^6^A depletion promoted partiality toward neuroectoderm at the expense of germline and mesodermal differentiation (Supplementary Fig. 7k-m).

To assess potential relationships between m^6^A deposition on transcripts and differentiation partiality, we characterized transcriptome-wide m^6^A distributions in human (4i hPSC) and mouse (FS) formative cells. As previously reported^60,61^, both human and mouse cells displayed preferences for similar m^6^A deposition sequences (Supplementary Fig. 7n). Also, both cell types favoured m^6^A deposition at transcript UTR’s, CDS’s and stop codons over introns, intergenic and non-coding regions (Supplementary Fig. 7o). A comparison of 4i hPSC and mouse FS cell m^6^A methylomes by meRIP-seq showed 1,983 human and 2,368 mouse m^6^A modified transcripts, with an overlap of 1,320 (30%) (Fig. 7j).

A recent study also showed that in iMeLCs, m^6^A abundances promote IGF2BP1 binding on the 3’UTR of *OTX2* transcripts, stabilizing them and preventing germline entry^28^. By contrast, we found no significant enrichment of m^6^A reads over input on human *OTX2* mRNA in 4i hPSCs. This is consistent with the absence of changes in m^6^A levels upon METTL3 inhibition by STM2457 treatment (Fig. 7k). This also coincides with the absence of *OTX2* transcript stability effects upon m^6^A depletion in 4i hPSCs (Supplementary Fig. 7c). In contrast, mouse *Otx2* showed enrichment of m^6^A reads over input and correspondingly, a significant reduction of m^6^A levels upon STM2457 treatment (Fig. 7k). We confirmed these findings by meRIP-qPCR, highlighting the absence of m^6^A abundance regulation on the 3’UTR of human *OTX2* transcripts (Fig. 7l). This region was previously shown to undergo significant m^6^A deposition and m^6^A level dependent transcript regulation in iMeLCs^28^.

Gene ontology (GO) analysis of m^6^A modified mRNAs in human (4i hPSCs) and mouse (FS) cells revealed preferential, lineage dependent m^6^A deposition. In mouse, m^6^A accumulation occurred predominantly on mesodermal transcripts (Fig. 7m). For example, *Mbd3*, a modulator of *Brachyury T* levels^62,63^ showed robust m^6^A deposition and STM2457 dependent m^6^A level regulation, unlike its human counterpart (Fig. 7n, o). This was associated with the upregulation of mouse, but not human *MBD3* expression in response to STM2457 treatment (Fig. 7p). In human, m^6^A accumulation predominated on neuroectodermal transcripts (Fig. 7m). Accordingly, *PAFAH1B1*, central to human neurodevelopmental migration^64^ showed high m^6^A deposition and STM2457 dependent m^6^A level regulation, unlike its mouse homolog (Fig. 7q, r). This was associated with a significant upregulation of human, but not mouse *Pafah1b1* in response to STM2457 (Fig. 7s). By meRIP-seq and GO term analysis, we identify for the first time, preferential m^6^A deposition on human neuroectodermal and mouse mesodermal transcripts as a potentially central mechanism for efficient differentiation partiality. We associate this with observed increases in differentiation propensities toward human neuroectoderm and mouse mesoderm in m^6^A depleted formative cells.

## Discussion

We demonstrate the essential roles of m^6^A abundances in the establishment and maintenance of formative pluripotency. Depletion of m^6^A in formative cells alters signaling activation, epigenetic remodeling, and TF transcript stabilities, restricting germline entry capacity. In m^6^A depleted EpiLCs and FS cells, increased transcript stability and expression of *Eras*, a mouse embryo specific, constitutively active RAS-like GTPase, activates PI3K and downstream pAKT signaling^43^. In line with transcript-reader specificity over redundancy mechanisms^23^, *Eras* transcripts are degraded primarily by YTHDF2, but not YTHDF3. In the mouse embryo, *Eras* expression is associated with mouse pre- and post-implantation epiblast expansion and declines with the onset of mesodermal differentiation^46^. Proteomic analyses demonstrate that pAKT diminishes throughout EpiLC state traverse^65^ while as shown here, *Eras* upregulation maintains pAKT activation upon m^6^A depletion. A recent report indicates that sustained pAKT blocks nuclear entry of FOXO TFs, preventing the establishment of a formative pluripotency-specific gene regulatory network^31^.

By profiling *Oct4*ΔPE::GFP reporter cells, we identified that pAKT activation also facilitates the disruption of *Oct4* distal enhancer decommissioning. Distal to proximal *Oct4* enhancer switching appears to represent a global ‘resetting’ of the naïve mouse epigenome during germline competence acquisition^26,53,66^. Either H3K27me3 or H3K9me2/3 repressive histone marks execute enhancer silencing dynamics upon cell differentiation^29,54,67^. We found globally depleted H3K27 trimethylation and correspondingly, downregulation of histone-lysine N-methyltransferase EZH2, which deposits the H3K27me3 histone mark. Intriguingly, co-IP analysis shows that overexpressed ERAS protein can physically interact with EZH2 in EpiLCs, however only in an m^6^A depleted setting. In accordance, *Eras* overexpression in wild-type EpiLCs cannot phenocopy germline entry deficiency observed upon m^6^A depletion. Hence, decreasing m^6^A abundances appears to alter protein-protein interaction networks with potential for ERAS mediated EZH2 repression and pAKT dependent modulation.

ChIP-seq analysis of H3K27 trimethylation further shows that decreasing m^6^A levels in EpiLCs results in the loss of diffuse, low abundance H3K27 trimethylation at genomic regions. This accounts for the bulk of cellular H3K27me3 genome-wide^55^. Hypomethylation of H3K27me3 on mESC and EpiLC enhancers is accompanied by a modest hypermethylation of H3K27me3 on day 2 PGCLC enhancers. This correlates with the unscheduled upregulation of mESC and EpiLC associated genes and repression of early germline markers. Depletion of m^6^A therefore interrupts formative state gene expression and appears to interfere with responsiveness to germline entry. Reversion toward a hyper-pluripotent state^25,35^ however is unlikely as *Nanog* becomes rapidly downregulated in EpiLC cultures. Also, m^6^A depleted cells lose the capacity to re-activate naïve, distal enhancer mediated *Oct4* expression upon transfer to the naïve conditions. In line with this, m^6^A depletion during mouse FS cell induction beyond day 2, suggests that *Oct4* distal enhancer silencing is not suppressed but delayed.

Unlike m^6^A depleted primed pluripotent mEpiSCs which quickly become lost to spontaneous differentiation or death^25^, mouse FS cells can be propagated long-term. However, m^6^A depletion impairs germline entry and sensitizes the cells toward spontaneous mesendodermal differentiation. This is associated with the emergence of a cell population illegitimately co-expressing the *Blimp1*-GFP reporter as well as *Otx2, Eomes, T* and *Flk1* mesendodermal markers. Nearly all *Blimp1*-GFP positive cells lack germline competence and likely contribute to germline entry deficiency observed upon m^6^A depletion in FS cells. Conversely, mesodermal induction is significantly enhanced. *Otx2* is a repressor of germline entry and inhibits the expression of *Prdm14* and *Blimp1*^27^. However, *Blimp1* co-expression with *Otx2* in m^6^A depleted FS cells is a by-product of both ERAS-PI3K-pAKT upregulation and increased *Blimp1* transcript stability. *Eras* knock-down rescues germline entry deficiency found in m^6^A depleted FS cells and in wild-type counterparts, *Eras* overexpression can partially phenocopy the impairment in germline entry observed upon m^6^A depletion. Recently, *Esrrb* was reported to prevent the illegitimate expression of mesendodermal markers in formative cells^68^. Dynamic m^6^A abundances may coordinate FOXO and ESRRB TF levels, enabling formative pluripotency while preventing mesendodermal TF upregulation and differentiation partiality.

Correspondingly, m^6^A depletion in human germline competent 4i hPSCs impairs germline entry efficiency from 20 to 10%. This is accompanied by the illegitimate co-upregulation of both *OTX2* and germline genes *STELLA* and *DDX4*. Neuroectodermal differentiation partiality coincides with mesodermal differentiation deficiency in m^6^A depleted cells. Compared to mouse, this bias may reflect *OTX2* and *SOX11* co-expression, associated with the distinctly ectodermal nature of human epiblast progression^11,12,69,70^. A recent study contrasts with our findings, suggesting that m^6^A abundances promote IGF2BP1 binding on the 3’UTR of *OTX2* transcripts, stabilizing them and preventing germline entry^28^. Hence, decreasing m^6^A levels increases germ cell induction in iMeLC conditions from 6 to 12%^28^. These dissimilarities likely reflect alternate regulatory mechanisms due to the capture of different formative states or adaptation to varying culture conditions. Despite of an apparent relatedness between transcriptomes^13^, FGF addition was shown to be detrimental to iMeLC growth^13^ and essential for 4i hPSC maintenance^16,71^, highlighting functional differences. We find that in 4i hPSCs, *OTX2* transcript stability is unaffected by m^6^A depletion, unlike that observed for *STELLA* and *DDX4* mRNAs. Accordingly, meRIP-seq analysis revealed the absence of m^6^A enrichment on human *OTX2* transcripts. This is confirmed by meRIP-qPCR on the 3’UTR of OTX2 mRNA, a region where IGF2BP1 was shown to bind in iMeLCs^28^.

Hence, we propose a different m^6^A dependent mechanism of OTX2 regulation in germline competent 4i hPSCs. Here, increased *FGF5* and *FGF10* transcript stabilities are associated with pFGFR1 and pERK activation, promoting OTX2 expression. Either mechanism may be found in human formative cells *in vivo*. Especially, given models of a fluctuating m^6^A epi-transcriptome in developmental progression^17^ and a need for OTX2 upregulation to protect the soma from germline program activation^27^. Hence, we show for the first time how an OTX2 dependent cell intrinsic barrier to germline entry in formative cells, can remain during a decrease in m^6^A level dynamics. *OTX2* knock-down in m^6^A depleted 4i hPSCs rescues the downregulation of germline genes *SOX17, OCT4, KLF4* and *NANOS3* in nascent hPGCLC spheres. This is consistent with a germline repressive role of OTX2 in both mouse and human formative cells^27,28,59^. However, *OTX2* knock-down in an m^6^A depletion setting was unable to restore efficient germline entry. This is likely due to the persistent downregulation of germ cell associated TFs like *BLIMP1, PRDM14, TFAP2C* and *NANOG*. Perhaps m^6^A depletion promotes pERK and transcript stability dependent upregulation of developmental TFs other than OTX2, responsible for activating somatic cell lineage enhancers and the repression of germline genes^59,69^.

Finally, we observed preferential m^6^A decoration of mesodermal transcripts in mouse and neuroectodermal transcripts in human, associated with increased propensities for differentiation into these lineages upon m^6^A depletion. Mechanisms governing m^6^A deposition on transcripts belonging to a specific cell lineage, remain an intriguing subject for further study. In future, perhaps timed m^6^A abundance manipulation can facilitate more efficient lineage induction and potentially, tissue maturation protocols from hPSCs. As in murine and porcine systems, only a fraction of wild-type human epiblast cells commits to the germ cell lineage^72-74^. Further exploration of m^6^A dependent mechanisms enabling homogeneous human germline entry, will aid in the treatment of infertility and germ cell cancer.

## Methods

### Cell Lines

Mouse cell lines *Oct4*ΔPE::GFP, *Blimp1*-GFP, *Nanog*VENUS, R1 and human cell lines H1, H9 and *OTX2*-YFP were cultured in humidified incubators at 37°C in 5 % CO_2_. Cells were routinely examined for the absence of mycoplasma contamination using the Mycoplasma Detection Kit.

### mESC culture in 2i+LIF

2i+LIF naïve mESCs were maintained inN2B27-based media as described previously^57^. Briefly, 500 ml of N2B27 contains 240 ml DMEM/F12, 240 ml Neurobasal, 5 ml N-2 supplement, 10 ml B-27 supplement, 1× Nonessential Amino Acids (NEAA), 1 mM sodium pyruvate, 0.1 mM β-mercaptoethanol, 1× Penicillin-Streptomycin-Glutamine. mESCs were cultured on Fibronectin (10 μg/ml) coated plate/flask in N2B27-based medium with 1000 U/ml mouse Leukemia Inhibitory Factor (LIF), 1 μM MEK inhibitor PD0325901 and 3.3 μM GSK3 inhibitor CHIR99021. Colonies were passaged by dissociating with TrypLE and the seeding density is 1.2 × 10^4^/cm^2^. ESCs originally cultured in FBS containing medium were adapted to N2B27 based 2i+LIF medium for a minimum of five passages.

### Induction of EpiLCs and pharmacological treatments

EpiLCs were induced as described previously^4^. Briefly, mESCs were seeded at 3 × 10^4^/cm^2^ on Fibronectin-coated plates in N2B27 medium containing 20 ng/ml Activin A, 12 ng/ml basic Fibroblast Growth Factor (bFGF), and 1% Knockout Serum Replacement (KSR). The medium was changed every day. In the case of chemical treatments (Supplementary Fig. 2g), 3 hours after seeding, EpiLCs were incubated with 10 μM STM2457, 5 μM LY294002 or 1 μM MK-2206 to inhibit METTL3, PI3K, or AKT respectively. Inhibitors were reconstituted in Dimethyl Sulfoxide (DMSO), applied at the same concentrations in control and treated conditions. The concentration of STM2457 is based on the IC_50_ value for STM2457 binding to mouse METTL3 (2.81 μM)^38^. Dose-response analysis of STM2457 is performed in EpiLCs (Supplementary Fig. 1 k, n). The concentration of LY294002 or MK-2206 is also based on a dose-response analysis (Fig. 3c and Supplementary Fig. 3e or 3f). Re-introduction of EpiLCs to 2i+LIF was performed as described previously^26^. Day 1 or day 2 EpiLCs were dissociated with TrypLE and seeded (1.2 × 10^4^/cm^2^) on Fibronectin-coated plates in N2B27 medium containing 2i+LIF for 3 days. The medium was changed daily.

Mouse formative stem (FS) cell culture. Mouse FS cells were obtained from wild-type R1 or *Blimp1*-GFP reporter line ESCs in 2i+LIF conditions by culturing in A_lo_XR medium^2^ for at least 10 days (Supplementary Fig. 5d). Briefly, A_lo_XR media contains Activin A (3 ng/ml), WNT agonist XAV939 (2 μM), and Retinoic Acid Receptor inhibitor BMS493 (1 μM) in N2B27 basal medium. FS cell clumps were dissociated with Accutase and passaged on 16.7 μg/ml Fibronectin–coated plates every 2 to 3 days. Medium was replaced daily. Chemical treatments occurred 3 hours after seeding. FS cells were incubated with DMSO control, 5 μM LY294002 or 5 μM STM2457. Dose-response analysis of STM2457 is also examined in FS cells (Supplementary Fig. 5k and 6a).

### Mouse fPSC culture

fPSCs were maintained as previously described^9^ in the Oct4ΔPE::GFP line ESCs. Briefly, 1 million cells dissociated in 100 μl N2B27 medium was incubated with 100 μl Matrigel at 37 °C for 1 hour for solidification. The cells embedded in Matrigel were filled with pre-warmed N2B27 medium supplemented with 1% KSR, 20 ng/ml Activin A,12 ng/ml bFGF and 5 μM XAV939, and cultured at 37° C and 5% CO_2_ for 3 days. For PGCLC induction, fPSCs were dissociated to single cells with TrypLE for 1 min at 37 °C.

### Human 4i culture

Human H1 and H9 cell lines were initially cultured on Mouse Embryonic Fibroblast (MEF) feeder cells in conventional hPSC conditions; DMEM/F12 containing 20% KSR, 1× NEAA, 1 mM sodium pyruvate, 0.1 mM β-mercaptoethanol, 1× Penicillin-Streptomycin-Glutamine and 4 ng/ml human FGF2. The *OTX2*-YFP (AI11e-*OTX2*YFP) cell line was initially cultured in mTeSR feeder free conventional hPSC conditions on Matrigel matrix. hPSCs were adapted for at least 2 weeks to 4i conditions; Knockout DMEM containing 20% KSR, 1 × NEAA, 1 mM sodium pyruvate, 0.1 mM β-mercaptoethanol, 1× Penicillin-Streptomycin-Glutamine, 8 ng/ml human FGF2, 20 ng/ml human LIF, 1 ng/ml human Transforming Growth Factor-β1 (TGF-β1), 1 μM PD0325901, 3 μM CHIR99021, 5 μM p38 inhibitor SB203580 and 5 μM JNK inhibitor SP600125^14^. Every 4 to 6 days, 4i hPSCs were dissociated with TypLE at passage ratio between 1:3 and 1:6. Culture medium for the first day after passage contained 10 µM ROCK inhibitor Y-27632. Medium was changed daily. Chemical treatments (DMSO control, 5 μM STM2457, 5 μM LY294002, 2 μM PD0325901 or 100 nM FGFR1 inhibitor PD173074) were conducted 1 day after seeding, and cells were collected 3 days after treatment.

### PGCLC induction

PGCLCs were induced in Nunclon Sphera 96 Well Round Bottom Low-Attachment Plate wells and GK15 medium with cytokines. GK15 medium contains GMEM, 15% KSR, 1 × NEAA, 1 mM sodium pyruvate and 0.1 mM β-mercaptoethanol. Cytokines include 500 ng/ml of Bone Morphogenetic Protein 4 (BMP4), 100 ng/ml of Stem Cell Factor (SCF) and 50 ng/ml of Epidermal Growth Factor (EGF). Mouse PGCLC medium contained 1000 U/ml mouse LIF, while human PGCLC medium contained 100 ng/ml human LIF. 10 µM Y-27632 and 0.25% (w/v) Poly (vinyl alcohol) were added to mouse fPSC, FS cell and human media to improve sphere formation^75^. Spheres cultured in GK15 medium without cytokines served as negative controls for flow cytometry analyses. Cell densities for seeding in each 96 well were 1 × 10^3^, 3 × 10^3^ and 4 × 10^3^ for mouse EpiLCs, FS cells and human 4i cells, respectively.

### Somatic cell lineage differentiation

Somatic cell lineage differentiations into mesoderm, endoderm, and ectoderm for mouse and human cells were performed as described previously^2^.

For human and mouse mesodermal induction, 3 × 10^3^/cm^2^ cells were seeded on 10 μg/ml Fibronectin-coated plates. Mouse FS cells were seeded in N2B27 basal medium containing 20 ng/ml Activin A and 3 μM CHIR99021 for 2 days. hPSCs were seeded in N2B27 basal medium containing 3μM CHIR99021 and 500 nM LDN193189 for the first 2 days followed by the addition of 20 ng/ml of human FGF2 for a further 2 days.

For human definitive endodermal induction, 6 × 10^3^/cm^2^ hPSCs were seeded on 10 μg/ml Fibronectin-coated plates (overnight at 4⍰°C) in N2B27 basal medium containing 100 ng/ml Activin A, 100 nM PI-103, 3 μM CHIR99021, 10 ng/ml human FGF2, 3 ng/ml BMP4 and 10 μg/ml Heparin for the first 24 hours and then replaced with 100 ng/ml Activin A, 100nM PI-103, 20 ng/ml FGF2, 250 nM LDN193189 and 10 μg/ml Heparin for a further 2 days.

For human neuroectodermal induction, 3 × 10^3^/cm^2^ hPSCs were seeded on the plates (percolated with 10 μg/ml Fibronectin and 10 μg/ml Laminin overnight at 4⍰°C) in N2B27 basal medium containing 1 μM A83-01 and 500 nM LDN193189 for 4 days. Medium was changed every second day.

### siRNA transfection

Lipofectamine RNAiMAX was used according to the manufacturer’s protocol. siRNAs we were purchased from Invitrogen. Mouse siRNAs were transfected into mouse EpiLCs (Supplementary Fig. 2m) or FS cells 3 hours after plating, and cells were collected 2 days after transfection. Transfection of human *OTX2* siRNA was performed in the 4i hPSCs one day after seeding, and cells were collected 3 days after transfection. For each gene, a mixture of two siRNA oligos was transfected. For *Akt* knock-down, a mixture of 6 siRNA oligos was transfected to simultaneously knock-down *Akt1, Akt2* and *Akt3*. For double knock-down of *Ythdf1, Ythdf2* or *Ythdf3*, a mixture of 4 siRNA oligos was transfected. The final working concentration of each siRNA mixture was 100 nM. The siRNA knock-down efficiency was examined by qPCR or western blots.

### Viral transduction

Lentiviral constructs containing iRFP fluorescent protein, Puromycin or Zeocin resistance and shRNAs sequences were self-designed and constructed by Vector Builder. 3 hours after cell plating, viral particles were added to cell supernatant at 75 MOI. After 2 days, cells were selected by 0.5 µg/ml Puromycin or 3 µg/ml Zeocin to achieve stable gene knock-down (Supplementary Fig. 2m, 5a and 5d). Transduction and selection efficiencies were monitored by flow cytometric iRFP detection. shRNA sequences were obtained from the Broad Institute’s The RNA interference (RNAi) Consortium. The lentiviral vectors over-expressing the control stuffer sequence or *Eras* sequence (5 MOI) were transduced in d0 EpiLC or FS cells.

### Real-Time quantitative PCR (RT-qPCR)

Total RNAs, free of genomic DNA, were extracted by Quick-RNA Microprep Kit and reverse transcribed by using PrimeScript RT Master Mix kit. Over 20 day 4 PGCLC spheres were used to obtain sufficient RNA yield. qPCR was carried out by using PowerUp SYBR Green Master Mix on a StepOnePlus Real-Time PCR System. *Rpl7* or *GAPDH* served as housekeeping genes for quantifying the relative mRNA abundances using the 2^−ΔΔCt^ method. qPCR primers are listed in Supplementary Table 1.

### m^6^A dot blot assay

Total RNA was isolated using TRIzol Reagent. Genomic DNA was removed by RQ1 RNase-Free DNase. Polyadenylated mRNA was purified using the Dynabeads mRNA Purification Kit. Purified mRNA was used for m^6^A dot blot as described previously^57^. In brief, 400 ng denatured mRNA was applied to rehydrated Hybond-N^+^ membrane within Bio-Dot Apparatus by gentle suction vacuum. The membrane was then cross-linked (UV, 5 minutes) and dried (801°C, 30 minutes) followed by blocking with 5% non-fat milk in Tris-buffered saline with 0.1% Tween (TBST) and then incubated with rabbit anti-m^6^A antibody (1/1000) (4⍰°C, overnight) and secondary incubation with Donkey Anti-Rabbit HRP antibody. Blots were developed by ECL reagents and scanned by Bio-Rad ChemiDoc XRS+ system. Methylene blue staining was used as RNA loading control.

### Mass spectrometry and proteomic data analysis

mRNAs were extracted three times by using Dynabeads™ mRNA Purification Kit from the total RNAs, which were extracted as previous described. The purities of mRNAs were confirmed by HS RNA screen tape. The quantification of RNA modification in mRNA samples was analyzed by Proteomics and Modomics Experimental Core Facility (PROMEC) at the Norwegian University of Science and Technology.

### Colorimetric m^6^A quantification

Total RNAs, free of genomic DNA, were extracted by Quick-RNA Kit as described above. The EpiQuik m^6^A RNA Methylation Quantification Kit was applied according to manufacturer’s instructions. The m^6^A signal in each sample was read to OD_450_ intensity on a micro plate spectrophotometer. Quantification of m^6^A levels in total RNA was calculated using a standard linear regression curve.

### Western blots

Total proteins were extracted by RIPA lysis buffer, containing Halt Protease Inhibitor Cocktail. For detecting global changes of histone marks, Histone Extraction Kit was applied. Protein concentration was quantified with Bradford assay. Protein samples were denatured (70 °C, 10 minutes) with the loading and reducing buffers. Electrophoresis and blotting were performed as described previously^57^. Blots were developed by ECL reagents and scanned using Bio-rad ChemiDoc XRS+ system. Histone H3 was used as a protein loading control for samples isolated by the Histone Extraction Kit. GAPDH was used as a protein loading control for samples isolated by RIPA lysis buffer.

### Immunofluorescence staining

Cells were grown on chambered, removable coverslips (Ibidi) coated with Fibronectin, and fixed with 4% paraformaldehyde at room temperature for 5 minutes. After permeabilization in phosphate-buffered saline (PBS) containing 0.25% Triton X-100 (room temperature, 20 minutes) and blocking in PBS containing 5% donkey serum (room temperature, 1 hour), coverslips were incubated with primary antibodies in PBS containing 0.5% Bovine Serum Albumin (BSA) (4⍰°C, overnight). Next day, coverslips were washed with PBS and incubated with the secondary antibody combination. Negative control staining was performed using only secondary antibodies. After 3 washes, coverslips were counterstained with antifade mounting medium containing DAPI. Images of stained cells were then acquired by wide-field microscopy. The filter sets included: Filter Set 38 HE (Zeiss, 489038-9901-000) for the 488 nm channel, Filter set 43 HE DsRed (AHF, F36-508) for the 555-nm channel, Filter set Cy5 ET (AHF, F46-006) for the 647-nm channel, and Filter Set 49 (Zeiss, 488049–9901-000) for the DAPI channel.

### Live cell multi-colour flow cytometry

Mouse ESCs, EpiLCs (day 2), human PSCs and PGCLC spheres (day 4) were dissociated into single cells with TrypLE. Mouse FS cells were dissociated by Accutase. Dissociation was neutralized in cold PBS wash buffer with 0.5 % BSA. After mesodermal induction, mouse cells were dissociated by cell dissociation buffer, then neutralized in cold HANK’s wash buffer without Ca^2+^ and Mg^2+^, supplemented with 1% BSA. Immunolabeling was conducted in cold washing buffer (4⍰°C, 0.5 hour). Large clumps of cells were removed using a cell strainer (BD Biosciences). Cells were washed once before BD LSR Fortessa analysis. Blue 488 nm laser and 530/30 channel were used to analyse GFP or YFP fluorescence. Green 561 nm laser and 585/15 channel were used to analyse PE fluorescence. Violet 405 nm laser and 450/50 channel were used to analyse BV421 fluorescence. Red 640 nm laser and 670/14 channel were used to analyse Alexa Fluor 647, APC or eFluor 660 fluorescence. Red laser and 730/45 channel were used to analyse iRFP fluorescence. Negative signal intensity for each fluorescence signals was set to 10^3^. The data was further analysed on the FlowJo software. Fluorescence compensation was performed where required according to single stained controls.

### Fluorescence-activated cell sorting (FACS)

Day 2 *Oct4*ΔPE::GFP reporter line EpiLCs were sorted on a BD FacsAria III cell sorter with a BD FacsDiva software. Mouse *Blimp1*-GFP reporter line FS cells were sorted on a FACSMelody cell sorter with a BD FACSChorus software using a 100 µm nozzle. Sorting was conducted using “Yield” mode to enrich target populations. The purity of each sorted population was confirmed by flow cytometric analysis (Supplementary Fig. 1p and Supplementary Fig. 6j), RT-qPCR (Supplementary Fig. 6i) or western blotting (Supplementary Fig. 6k).

### High-throughput data sourcing

Refseq gene annotations^76^ for mm10 were downloaded from the refflat table at UCSC^77^ and imported as a Geneset into EaSeq. Gene subsets and putative enhancer data were acquired from Kurimoto et al ^29^ in their provided Supplemental Tables S3 and coordinates were converted from mm9 to mm10 using the liftover tool at UCSC^78^. ATAC-seq in mouse ESCs and FS cells was downloaded from Gene Expression Omnibus (GEO): GSE131556^2^. ATAC-seq broad peaks on distal and proximal enhancer of mouse *Oct4* was visualized on USCS Genome browser to the mouse reference genome (GRCm38/mm10).

### Transcriptomic profiling and data analysis

Total RNAs from shRNA transduced EpiLCs were extracted using TRIzol as described previously^57^. mRNA was purified by using poly-T oligo-attached magnetic beads. Following fragmentation, the RNA fragments were converted to first strand cDNA using reverse transcriptase and random primers. Second strand cDNA synthesis was conducted using DNA polymerase I and RNase H. The products were then enriched with PCR amplification. The PCR yield was quantified by Qubit and pooled together into a library of single stranded DNA (ssDNA) circles. DNA nanoballs (DNBs) were generated with the ssDNA circle by rolling circle replication (RCR) to enlarge the fluorescent signals. The DNBs were loaded into the patterned nanoarrays and pair-end reads of 100 bp were generated on the DNBseq platform. Low-quality reads, reads with adaptors and reads with unknown bases were filtered out. Mapping of clean reads was performed on the reference genome (mm9_UCSC_20180118) using HISAT2 (v2.0.4)^79^. RNA sequencing was measured in day 2 EpiLCs with three conditions, Scr, sh*Mettl3*-1 and sh*Mettl14*-1 from three independent experimental batches, resulting in nine RNA-seq samples. Fragments Per Kilobase of transcript per Million mapped reads (FPKM) was used for expression level normalization. DEseq2 was employed to detect differentially expressed genes (DEGs) in each batch by setting absolute log_2_Fold Change ≥1 and adjust *p* value (*p*_*adj*_ ≤0.05). Heatmaps, Venn plots and MA plots were generated using R (https://www.R-project.org) with the package pheatmap. The DEGs between Scr and sh*Mettl3*-1 are listed in Supplementary Table 3, and the DEGs between Scr and sh*Mettl14*-1 are listed in Supplementary Table 4.

Total RNA in *Blimp1*-GFP EpiLCs with DMSO or 10 μM STM treatment was extracted by Quick-RNA Microprep Kit. rRNA was removed by Next® rRNA Depletion Kit v2 (Human/Mouse/Rat), following the manufacturer’s protocol. The RNA library was prepared by using SMART-Seq Total RNA Mid Input kit, following the manufacturer’s protocol. After examination via HS DNA tape station, the RNA libraries were sequenced by the Norwegian Sequencing Centre on an Illumina NovaSeq X platform using PE150 read length. Low-quality reads, reads with adaptors, unmapped reads, rRNA-mapped reads and PCR duplicates were filtered out. Clean reads were aligned to the reference genome mm10 using STAR (v2.7)^80^. Values were normalized to FPKM. Gene expression quantification was derived from STAR-aligned reads. Differentially expressed genes (DEGs) between control and STM-treated samples were identified using EdgeR, applying the glmQLFTest function while accounting for batch effects. The DEGs between control (Ctr) and STM treatment are listed in Supplementary Table 5.

### Methylated mRNA-immunoprecipitation (meRIP) assay

Purified mRNA samples were chemically fragmented into 100-500 nucleotide fragments by incubation with RNA Fragmentation Reagents (751°C, 3.5 minutes). Fragmented mRNA was purified by using RNA Clean & Concentrator-5 kit and examined by Tape Station 4150 (Agilent). 50 ng of mRNA fragments were left as input, while 1 μg of mRNA fragments were immunoprecipitated (IP) with 2.5 μg mouse anti-m^6^A antibody or IgG in 250 μl IPP buffer (10 mM Tris-HCl, pH 7.4, 150 mM NaCl and 0.1% NP-40) (4⍰°C, overnight). Next day, 750 μg Dynabeads Protein G were added to pull down m^6^A-modified mRNA fragments (4⍰°C, 5 hours). After 3 washes with ice-cold IPP buffer, bound m^6^A-methylated RNA fragments were eluted with 0.5 mg/ml N^6^-Methyladenosine (25⍰°C, 60 minutes) and then purified by RNA Clean & Concentrator-5 kit. All the RNA samples from IP and input were reverse transcribed with High-Capacity cDNA Reverse Transcription Kit. The m^6^A binding sites on *Eras* and *Blimp1* mRNAs were predicted on Primer Premier 5 based on the known RRACH m^6^A motif sequence^19^ (Supplementary Fig. 2d and 6h). Primers are listed in Supplementary Table 1. Enrichment was calculated by the formulation of % Input 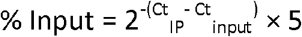

### Methylated RNA immunoprecipitation sequencing

Methylated RNA immunoprecipitation sequencing (MeRIP-seq) was conducted as previously described ^60^. Briefly, total RNAs from mouse FS cells and human 4i hPSCs were extracted, and rRNA was removed by using Next® rRNA Depletion Kit v2 (Human/Mouse/Rat), following the manufacturer’s protocol. Then RNA samples were purified using 2.2× volume of RNAClean XP beads, washed twice with 80% ethanol, and eluted with nuclease-free water. RiboLock RNase inhibitor (1 U/µl) was added to each RNA sample to prevent RNA degradation. Approximately 200 ng rRNA-removed RNAs were sonicated by UP100H Ultrasonic Processor (Hielscher) equipped with a 2-mm probe. The samples underwent 2⍰× ⍰30⍰seconds sonication cycles, precisely alternating between 30⍰seconds of sonication and 30⍰seconds on ice for each cycle. The RNA fragments were examined by the HS RNA screen tape to ensure the main range is between 100 and 500-nucleotide-long. On the other hand, 10 μg of rabbit m^6^A antibody was conjugated with 750 μg Protein A Dynabeads in the IPP buffer at 4 °C overnight. After washing the antibody-bead with IPP buffer, about 100 ng RNA fragments were incubated with 1/10 volume of the antibody-beads in 100 μl IPP solution for immunoprecipitation at 4°C overnight, while about 50 ng of the fragmented RNA was reserved as input RNA. The RNA–antibody–bead complexes were washed four times on ice in the following order: 1, medium-stringency RIPA buffer (10⍰mM Tris-HCl (pH 8.0), 300⍰mM NaCl, 1⍰mM EDTA, 0.5⍰mM EGTA, 1% (vol/vol) Triton X-100, 0.2% (vol/vol) SDS and 0.1% (vol/vol) sodium deoxycholate); 2, high-stringency RIPA buffer (10⍰mM Tris-HCl (pH 8.0), 350⍰mM NaCl, 1⍰mM EDTA, 0.5⍰mM EGTA, 1% (vol/vol) Triton X-100, 0.23% (vol/vol) SDS and 0.1% (vol/vol) sodium deoxycholate); 3, medium-stringency RIPA buffer; 4, IPP buffer. The banded RNAs were eluted twice at 55⍰°C using a thermomixer at 12000 rpm in 150 μl buffer of 5⍰mM Tris-HCl (pH 7.5), 1⍰mM EDTA buffer, 0.05% (vol/vol) SDS, Proteinase K (11.2 U/ml) and RiboLock RNase inhibitor. The first-round elution lasted for 90 minutes and the second one lasted for 5 minutes. Proteinase K was then inactivated by placing the samples in 80⍰°C for 15⍰min after each elution. The eluted m^6^A antibody-binding RNAs and input RNAs were precipitated by adding 1/10 volume of 3M sodium acetate (pH 5.2), 50 mg linear acrylamide and 2.5× volume of cold ethanol and placed at −80 °C for 1 hr. After washing with cold 75% ethanol, the RNA samples were resuspended in water and examined by HS RNA screen tape for library preparation.

The RNA libraries for both the input and IP samples were constructed using SMART-Seq Total RNA Mid Input kit, following the manufacturer’s protocol. After examination via HS DNA tape station, the RNA libraries were sequenced by the Norwegian Sequencing Centre on an Illumina NovaSeq X platform using PE150 read length.

### MeRIP-seq Analysis

MeRIP-seq analysis was carried out based on the MeRipBox pipeline (https://github.com/Augroup/MeRipBox) with several modifications. Raw reads were first assessed for quality using FastQC (v0.11.8). Adapter sequences and low-quality bases were trimmed using bbduk with the parameters: ktrim=r, k=23, mink=11, hdist=1, tbo, tpe, qtrim=r, trimq=15, maq=15, minlen=36, forcetrimright=149, threads=16. Trimmed reads were aligned to the mm10 (mouse) or hg38 (human) reference genome using STAR (v2.7)^80^ with the following options: -- outFilterMultimapNmax 1 and --quantMode GeneCounts, to retain only uniquely aligned reads and to enable gene-level quantification. SAMtools (v1.19)^81^ was used to remove PCR duplicates. Reads aligning to rRNAs were filtered out using BEDTools (v2.28.0)^82^, based on GENCODE vM23 for mouse and GENCODE v47 for human. Clean reads were used for m^6^A peak calling.

For visualization, BigWig files representing genome-wide coverage were generated using deepTools (v3.2.0)^83^ with bamCoverage (--binSize 10 --normalizeUsing RPKM) and visualized using IGV (v2.18.4). To assess m^6^A signal distribution, for each 10-bp bin, the signal was calculated as: log2((RPKM_IP + 1) / (RPKM_Input + 1)).

m^6^A peak calling was performed using MACS3 (v2.1.2)^84^ with the parameters -g 242010196 --keep-dup all -B --nomodel --call-summits for mouse samples and -g 300000000 --keep-dup all -B -- nomodel --call-summits for human samples. Peaks with a q-value < 0.05 were retained. A gene was defined as m^6^A positive if at least one transcript from the gene overlapped with ≥1 m^6^A peak in at least two biological replicates within exonic regions. Peak annotation was performed using BEDTools^82^ with GENCODE annotations (vM23 for mouse and v47 for human), using only the genomic coordinate of the peak summits (1 bp). If the summit of an m^6^A peak overlapped across multiple genomic features, it was assigned to a single feature based on the following priority: stop codon (±200 bp), 3’ UTR, 5’ UTR, CDS, intron, and intergenic. Metagene profiles of m^6^A peak summits across the 5’ UTR, CDS, and 3’ UTR of protein-coding genes were generated using MetaPlot^85^, following the recommended guidelines.

Motif enrichment analysis within m^6^A peak regions (±200 bp from the summit) was carried out using Homer (v4.11.1)^86^ with the parameters -rna -len 5,6,7,8.

For cross-species comparison, orthologous gene sets comprising 16,473 protein-coding genes were obtained from MGI (http://www.informatics.jax.org/), and only these genes were used in correlation analyses of m^6^A sites between human and mouse. Gene Ontology (GO) enrichment analysis was conducted using clusterProfiler^87^.

### Chromatin immunoprecipitation (ChIP)

EpiLCs were fixed with 1% formaldehyde for 10 minutes at room temperature to cross-link proteins at chromatin DNA, which was terminated with 1 M glycine. After washing, cells were scraped off and lysed with 1% sodium dodecyl sulfate lysis buffer supplemented with protease inhibitor cocktail on ice. The lysate was sonicated to shear chromatin DNA into fragments between 200 and 1000 base pairs. DNA fragments were purified by using ChIP DNA Clean & Concentrator Kit and the size of fragments was examined by Tape Station. After precleaning with Protein A/G Magnetic Beads for 1 hour at 4 °C, sheared chromatin DNA was immunoprecipitated at 4 °C overnight with 5 μg of antibody. IgG was used as a negative control. The immuno-precipitate was incubated with magnetic beads for 3 hours, and beads were magnetically pulled down. After washing, reverse cross-linking was performed in 5 M NaCl at 65°C overnight. Samples were purified from RNA and protein by RNase A and proteinase K treatment, respectively. Then, DNA was extracted using ChIP DNA Clean & Concentrator Kit. Sheared DNA without antibody precipitation after reverse cross-linking was used for input controls. *Oct4* distal enhancer primer sequences were adapted from a previous study^34^ and listed in Supplementary Table 1.

### Chromatin immunoprecipitation sequencing (ChIP-seq)

Chromatin immunoprecipitation sequencing (ChIP-seq) was performed in *Blimp1*-GFP EpiLCs as described in the ChIP section with a minor modification consisting of the DNA-antibody-bead incubation. 10 μg of H3K27me3 antibody was conjugated with 750 μg protein A dynabeads in 100 μl ChIP buffer (10⍰mM Tris-HCl (pH 8.0), 140⍰mM NaCl, 1⍰mM EDTA, 0.5⍰mM EGTA, 1% (vol/vol) Triton X-100, 0.15% (vol/vol) SDS and 0.1% (vol/vol) sodium deoxycholate, 1 mM PMSF, 1x protease inhibitor cocktail) at 4 °C overnight. Then the complex was washed twice with ChIP buffer before immunoprecipitation. About 20 μg chromatin DNA was incubated with the prewashed antibody-bead in 1 ml ChIP buffer at 4°C overnight. The immunoprecipitated chromatin DNA was examined by HS DNA Tape station and Qubit before sequencing. The library quality was assessed using Qubit for quantification, and size distribution was detected using a Bioanalyzer. Sequencing was performed on an Illumina Novaseq using PE150 read length by Shenzhen Acegen Technology Co., LTD (China).

### ChIP-seq data processing

Paired-end FASTQ files were processed using the ew-qctrimalign Nextflow pipeline (https://github.com/lerdruplab/ew-qctrimalign) with the --aligner bowtie2 option. Adapter and quality trimming were performed with Trim Galore v0.6.7 (https://github.com/FelixKrueger/TrimGalore). The trimmed reads were aligned to the Mus musculus mm10 reference genome using Bowtie2 v2.5.2^88^ in end-to-end alignment mode. Resulting BAM files were sorted using SAMtools v1.2^81^, converted to BED format with BEDTools v2.31.1^82^, and imported into EaSeq^89^ for duplicate removal and downstream analysis. Unless stated, all ChIP-seq values are normalized to FPKM.

### ChIP-seq data visualization and handling

H3K27me3 signal was quantified using the Quantify tool in EaSeq using standard FPKM normalization in the indicated span of the 10 kbp bins for MA-plots or at span from 2 kbp upstream to 2 kbp downstream the center of enhancers. Means and log_2_-fold differences were calculated using the Calculate tool in EaSeq. Genome browser tracks of H3K27me3 levels were plotted using the FillTrack plot type in EaSeq, while simple heatmaps of fold differences and gene overlap were made using the ParMap plot type. Heatmaps and ratiomaps with spatial resolution were made using the Heatmap and Ratiomap types, respectively. For H3K27me3 heatmaps of genes with significantly altered expression, genes with high levels of signal (Mean gene body input FPKM > 1) were excluded. MA-plots and colored MA-plots were based on mean H3K27me3 and log_2_ fold differences calculated as described above and visualized using the Scatter and Z-Scatter plot types in EaSeq. Identification of the nearest gene for each putative enhancer was done using the Coloc. tool in EaSeq set to use border to border distances.

### Bubble plots of gene overlaps

For bubble plots (Fig. 4m) of gene set overlaps and significance, first each gene was categorized as up/down/no-change using the log2fold change and an adjusted significance threshold of 0.05. Cell type expression categories were acquired from Kurimoto et al^29^. The log_2_ fold difference is calculated based on the observed number of genes in each combination of categories compared to the expected number, which was based on the global average frequencies. Statistical significance was assessed using Chi-square testing and p-values were Benjamini-Hochberg adjusted for multiple testing^90^. Code for the overall workflow can be found at (https://github.com/lerdruplab/m6A).

### mRNA degradation assay

5 μg/ml Actinomycin-D was added to mouse culture medium for 6, 3 and 0 hours before collection. The same concentration of Actinomycin-D was added to human culture medium 4 hours before collection. RNA degradation is calculated as described previously^57^. R^2^ is the correlation coefficient.

### Co-Immunoprecipitation (Co-IP)

The 5 μg of mouse ERAS antibody was conjugated with 750 μg protein G dynabeads in 100 μl RIPA lysis buffer overnight at 4 °C. The mouse IgG1 antibody was also conjugated with protein G dynabeads, respectively in the same condition as negative controls. About 10 million EpiLCs were lysed in 400 μl RIPA lysis buffer containing a protease inhibitor cocktail for at least 1 hour and centrifuged at 8000g x 10 min at 4°C. The supernatant was collected for immunoprecipitation and 5% of the supernatant was kept as input. The extract was incubated with prewashed anti-ERAS conjugated dynabeads on a rotating wheel at 4°C overnight. Beads were washed in cold PBS containing the protease inhibitor cocktail for 3 times and eluted by boiling in RIPA-diluted NuPAGE™ LDS Sample Buffer and Reducing Buffer at 70 °C for 10 min. The eluted proteins and input were analyzed by western blot. A-RAF and GAPDH was detected as positive and negative control for ERAS binding, respectively.

## Statistics and reproducibility

At least three independent experiments were performed for each assay. Data is presented as the mean ± standard deviation (SD). Statistical significance was assessed by two-tailed, paired t-test and listed in Supplementary Table 2. Non-significant (N.S.) differences are defined as *p* ≥0.1 by comparison. Immunoblot images are representative of at least three independent experiments. Uncropped versions are available in the Supplementary Files.

## Data and code availability

Raw high-throughput sequencing data have been deposited in the NCBI Sequence Read Archive (SRA). All data were analyzed with standard programs and packages, as detailed above. All raw images, immunoblots, and values for quantification are available from the corresponding author and lead contact on request.

## Authors’ contributions

R.Z. performed most experiments and analysed data with A.F. B.B. analyzed published ATAC-seq data^2^ and meRIP-seq experiments performed by R.Z. K.J. contributed to experimental technique development in EpiLCs. M.S. and E.I. analysed ChIP-seq and meRIP-seq experiments performed by R.Z. Y.Y. analysed RNA-seq experiments performed by R.Z. J.L. and L.W. assisted with cell preparations for assays. A.K., X.W and Y.L. contributed to meRIP-seq technique optimization. J.W. provided meRIP-seq analysis support. M.L. coordinated all analyses on ChIP-seq and most analyses on meRIP-seq experiments carried out by R.Z. A.F. designed, supervised, and acquired funding for the study. A.F wrote the manuscript with R.Z and M.L.

## Acknowledgments

We extend gratitude to Professor Azim Surani (University of Cambridge) for providing us with the *Oct4*ΔPE::GFP and *Blimp1*-GFP mESC reporter cell lines. We thank Professor Konstantinos Anastassiadis (BIOTEC, Technische Universitaet Dresden) for providing the *Nanog*VENUS mESC reporter cell line and together with Doctor Carsten Marr (Helmholtz Research Centre, Munich) and Professor Hilde Loge Nilsen (Oslo University Hospital), for offering critical review of the manuscript. We extend gratitude to Professor Renwei Su (South China Agricultural University) for supplying cell material for study and Doctor Cathrine Broberg Vågbø (PROMEC, Norwegian University of Science and Technology) for aiding us in Mass Spectrometry experiments. We thank the Norwegian research infrastructure services (NRIS)-SIGMA2 for providing large data storage and High-performance computing (HPC) for our Project (NN11110K).

## Availability of data

All data is included in this article [and in Supplementary Files].

## Ethics approval and consent to participate

Not Applicable.

## Consent for publication

Not Applicable.

---

**Supplementary Figure 1. Depletion of m^6^A Impairs Germline Entry.**

**a-i**, R1 wild-type, *Oct4*ΔPE::GFP, *Blimp1*-GFP or *Nanog*VENUS mESCs were transduced with scramble control (Scr), *Mettl3* (shM3-1 or shM3-2) or *Mettl14* (shM14-1 or shM14-2) shRNA, followed by EpiLC or fPSC induction and PGCLC specification. See also Methods for experimental details. Mean ± SD. Two-tailed, paired t-test: vs Scr, \*\*\**p* <0.001; \*\**p* <0.01; \**p* <0.05.

**a**, Morphology of *Blimp1*-GFP day 2 EpiLCs. Scale bar, 100 μm.

**b**, Immunoblots of METTL3 and METTL14 protein levels in *Blimp1*-GFP ESCs and day 2 EpiLCs.

**c**, m^6^A dot blots showing m^6^A levels in *Blimp1*-GFP day 2 EpiLCs. IB, immunoblotting.

**d**, Mass spectrometry quantification of m^6^A, m^7^G and m^6^Am level changes in the mRNAs of *Oct4*ΔPE::GFP day 2 EpiLCs. N =3.

**e**, Graph summarizing FACS analyses of the *Blimp1*-GFP reporter positive cells proportion in day 4 PGCLC spheres derived from EpiLCs. N =5. See Supplementary Files for representative FACS plots.

**f**, Graph summarizing FACS analyses for the percentage of SSEA1 and CD61 double positive cells in day 4 PGCLC spheres derived from Oct4ΔPE::GFP fPSCs. N =3. See Supplementary Files for representative FACS plots.

**g**, FACS analysis showing the percentage of *Nanog*VENUS reporter positive cells in day 2 EpiLCs. Dashed line indicates positive gating. Solid gray line represents VENUS signals in positive control.

**h**, Immunofluorescence staining for OCT4 and NANOG in R1 ESCs and day 2 EpiLCs.

**i**, Graphs summarizing FACS analyses measuring the percentage of *Oct4*ΔPE::GFP reporter positive cells obtained from (left) day 1 or (right) day 2 EpiLCs re-introduced into 2i+LIF.

**j-o**, Day 0 R1 or *Oct4*ΔPE::GFP reporter EpiLCs were either (**j, m**) transduced with scramble control (Scr), *Mettl3* (shM3-1) or *Mettl14* (shM14-1) shRNA, or (**k-l, n-o**) treated with DMSO or METTL3 inhibitor STM2457 (STM) for 2 days, followed by PGCLC specification. See also Methods for experiment details. Mean ± SD. Two-tailed, paired t-test: vs 0 or Ctr or Scr, \*\*\**p* <0.001; \*\**p* <0.01; \**p* <0.05; vs 5, ^#^*p* <0.05; vs 10, ^&&^*p* <0.01.

**j**, qPCR showing (Left) *Mettl3* or (right) *Mettl14* mRNA levels in *Oct4*ΔPE::GFP day 2 EpiLCs. N =3.

**k**, Colorimetric m^6^A quantification analysis in R1 day 2 EpiLCs. N =4.

**l**, Mass spectrometry quantification of m^6^A, m^7^G and m^6^Am changes in the mRNAs of day 2 *Oct4*ΔPE::GFP reporter EpiLCs. N =3.

**m-o**, Graphs summarizing FACS analyses measuring (**m-n**) the percentage of *Oct4*ΔPE::GFP reporter positive cells in day 2 EpiLCs and (**o**) the percentages of SSEA1 and CD61 double positive cells in R1 day 4 PGCLC spheres. N =3. See also Supplementary Files for representative FACS plots.

**p-r**, *Oct4*ΔPE::GFP reporter mESCs were transduced with scramble control (Scr), *Mettl3* (shM3-1) or *Mettl14* (shM14-1) shRNA before EpiLC induction. Day 2 EpiLCs were sorted into GFP positive and negative cell fractions for PGCLC specification. See also Methods for experiment details.

**p**, FACS analysis showing the percentages of *Oct4*ΔPE::GFP reporter positive cells in day 2 EpiLC before and after GFP sorting. Dashed line indicates positive gating.

**q**, Live imaging showing *Oct4*ΔPE::GFP reporter signals (merged with bright field) in day 4 PGCLC spheres. Scale bar, 100 μm.

**r**, Graph summarizing FACS analyses showing the percentages of SSEA1 and CD61 double positive cells in day 4 PGCLC spheres. Mean ± SD. N ≥3; two-tailed, paired t-test: vs Scr GFP +, \*\**p* <0.01; \**p* <0.05; vs shM3-1 GFP -, ^#^*p* <0.05; vs shM14-1 GFP -, ^&&^*p* <0.01; N.S., Non-significant. See also Figure 1f for representative FACS plots.

**Supplementary Figure 2. m^6^A Dependent *Eras* Transcript Degradation in EpiLCs Ensures Robust Germline Entry.**

**a-i**, *Oct4*ΔPE::GFP or *Blimp1*-GFP reporter mESCs were transduced with scramble control (Scr), *Mettl3* (shM3-1 or shM3-2) or *Mettl14* (shM14-1 or shM14-2) shRNA, followed by EpiLC induction (**a-e, g-i**). Non-transduced *Blimp1*-GFP d0 EpiLCs were treated with DMSO or METTL3 inhibitor STM2457 for 2 days (**f**). Mean ± SD. See also Methods for experiment details.

**a**, MA plots of RNA-seq analyses in day 2 *Oct4*ΔPE::GFP reporter EpiLCs showing (upper panel) scramble control (Scr) vs *Mettl3* (shM3-1) shRNA and (lower panel) scramble control vs *Mettl14* (shM14-1) shRNA. X axis represents value A (log_2_ transformed mean expression level). Y axis represents value M (log_2_ transformed fold change). Red dots or circles represent upregulated differentially expressed genes (DEGs). Blue dots or circles represent downregulated DEGs. Gray points or circles represent genes with non-significant changes (N.S.). The number of genes upregulated (Up), downregulated (Down) or non-significantly changed is shown next to the plots.

**b**, Heat map showing the expression of pluripotency or lineage genes from RNA-seq analyses in day 2 *Oct4*ΔPE::GFP reporter EpiLCs. The color represents log_10_ (FPKM). FPKM, Fragments Per Kilobase of transcript per Million mapped reads.

**c**, Immunoblots showing OTX2 expression in *Blimp1*-GFP ESCs and day 2 EpiLCs.

**d**, Diagram showing the putative m^6^A motif sites on *Eras* mRNA and the primers designed for meRIP-qPCR. UTR, untranslated region. CDS, coding sequence.

**e-g**, meRIP-qPCR in *Oct4*ΔPE::GFP reporter cells showing abundance of m^6^A on *Eras* mRNA in ESCs (**e**, N =4), EpiLCs (**f**, N =3). meRIP-qPCR in *Oct4*ΔPE::GFP reporter cells showing abundance of m^6^A on *Oct4* and *GFP* mRNA, compared to *Eras* mRNA in EpiLCs transduced with scramble control shRNA (**g**, N =3). Two-tailed, paired t-test: vs Scr or Ctr or *Eras*,, \*\*\**p* <0.001;*p <0.05.

**h-i** qPCR showing stabilities for *Oct4* (**h**) and *GFP* (**i**) mRNAs after Actinomycin-D treatment in day 2 *Oct4*ΔPE::GFP reporter EpiLCs. N =3. Statistical analysis details are listed in Supplementary Table 2.

**j-l**, Day 0 *Oct4*ΔPE::GFP reporter EpiLCs were transfected with negative control (Nc), *Ythdf1* (siY1), *Ythdf2* (siY2), *Ythdf3* (siY3) or *Ythdf1* and *Ythdf2* (siY1+2) siRNA for 2 days.

**j**, qPCR for *Eras* mRNA stability assay. N ≥4. Statistical analysis details are listed in Supplementary Table 2.

**k-l**, Immunoblots of YTHDF1, 2, 3 and ERAS expression and AKT phosphorylation in day 2 EpiLCs.

**m-q**, *Oct4*ΔPE::GFP or *Blimp1*-GFP reporter mESCs were transduced with scramble control (Scr), *Mettl3* (shM3-1 or shM3-2) or *Mettl14* (shM14-1 or shM14-2) shRNA, followed by EpiLC induction and PGCLC specification. See also Methods for experiment details. Mean ± SD. N =4; two-tailed, paired t-test: vs Scr Nc, \*\*\**p* <0.001; \**p* <0.05; vs shM3-1 Nc,^##^*p* <0.01;^#^*p* <0.05; vs shM14-1 Nc, ^*&&&*^*p* <0.001;^&^*p* <0.05; N.S., Non-significant.

**m**, Schemata illustrating experimental strategy. PD, MEK inhibitor PD0325901. CHIR, GSK3 inhibitor CHIR99021; LIF, Leukemia inhibitory factor. bFGF, basic Fibroblast Growth Factor. ActA, Activin A. BMP4, Bone Morphogenetic Protein 4. SCF, Stem Cell Factor. EGF, Epidermal Growth Factor.

**n-o**, qPCR showing *Eras* mRNA levels in day 2 *Oct4*ΔPE::GFP reporter EpiLCs (**n**) and day 4 PGCLC spheres (**o**).

**p-q**, Graph summarizing FACS analyses of day 4 *Blimp1*-GFP reporter PGCLC spheres showing either (**p**) percentages of SSEA1 and CD61 double positive cells, or (**q**) the proportion of *Blimp1*-GFP positive cells. See also Figure 2h and Supplementary Files for representative FACS plots.

**Supplementary Figure 3. Depletion of m^6^A and *Eras* Upregulation Impair Germline Entry by Reducing H3K27 Trimethylation.**

**a-i**, *Oct4*ΔPE::GFP reporter ESCs were transduced with scramble control (Scr), *Mettl3* (shM3-1) or *Mettl14* (shM14-1) shRNA before EpiLC induction. Day 0 EpiLCs were either transfected with negative control (Nc), *Eras* (si*Eras*), or *Akt* (si*Akt*) siRNA, or treated with DMSO or PI3K inhibitor LY294002 (LY) or AKT inhibitor MK-2206 (MK) for 2 days. See also Figure S2m and Methods for experimental protocol. Mean ± SD. Two-tailed, paired t-test: vs Scr Nc or Scr, \*\**p* <0.01; \**p* <0.05; vs shM3-1 Nc or shM3-1, ^#^*p* <0.05; vs shM14-1 Nc,^&^*p* <0.05; N.S., Non-significant.

**a-b**, Representative FACS plot showing the proportion of *Oct4*ΔPE::GFP reporter positive cells in day 2 EpiLCs transfected with Nc or *Eras* (**a**) or AKT (**b**) siRNA. Dashed line indicates positive gating.

**c**, Graph summarizing effects in (**b**). N ≥3.

**d-f**, Immunoblots of total AKT and phosphorylated AKT levels in day 2 EpiLCs transfected with Nc or *Akt* siRNA (**d**), or treated with control DMSO or LY294002 (LY) (**e**) or MK-2206 (**f**).

**g-h**, Graph summarizing FACS data measuring (**g**) either the proportion of *Oct4*ΔPE::GFP reporter positive cells upon DMSO or MK-2206 treatment in day 2 EpiLCs, (**h**) or the percentage of SSEA1 and CD61 double positive cells in day 4 PGCLC spheres derived from these treated EpiLCs. N =3. See also Supplementary Files for representative FACS plots.

**i**, Immunoblots of H3K27me3 levels in day 2 EpiLCs.

**j-l**, Day 0 *Oct4*ΔPE::GFP reporter EpiLCs were transduced with overexpression lentivirus for either control (OE ctr) or ERAS (OE Eras), and at day2 these EpiLCs were induced into PGCLC for 4 days.

**j**, Immunoblotting analysis of ERAS, AKT phosphorylation and H3K27me3 in day 2 EpiLCs.

**k-l**, Graph summarizing FACS data measuring either (**k**) the proportion of *Oct4*ΔPE::GFP reporter positive cells in day 2 EpiLCs, or (**l**) the percentage of SSEA1 and CD61 double positive cells in day 4 PGCLC spheres. N =3; two-tailed, paired t-test: N.S., Non-significant. See also Supplementary Files for representative FACS plots.

**m**, Day 0 *Blimp1*-GFP reporter EpiLCs were transduced with overexpression lentivirus for either control (OE ctr) or ERAS (OE Eras), followed by treatment with DMSO or the METTL3 inhibitor STM2457 (STM) for 2 days. Co-immunoprecipitation (Co-IP) was performed using an ERAS antibody, with subsequent immunoblotting probed for EZH2 (tested interaction), A-RAF (positive interaction control), GAPDH (negative interaction and loading control), and IgG (antibody control) are shown, with input lysates included for reference.

**Supplementary Figure 4. Depletion of m^6^A and Reduced H3K27 Trimethylation Deregulates EpiLC Gene Expression.**

**a-s**, Day 0 *Blimp1*-GFP reporter EpiLCs were treated with DMSO (Ctr) or METTL3 inhibitor STM2457 (STM) for 2 days, then collected for H3K27me3 ChIP sequencing (**a, c-s**) and RNA sequencing (**b**) analyses.

**a**, DNA Tape Station showing H3K27me3-enriched chromatin DNA in biological replicate 1 before library preparation.

**b**, Heatmap showing the log_2_ fold difference in expression for genes significantly affected upon STM treatment. See also Supplementary Table 5 for the details of gene expression changes.

**c-h**, Heatmaps showing the distribution of FPKM normalized H3K27me3 signal at the gene body and surrounding loci of 184 genes significantly affected upon STM treatment. Genes were ordered by expression difference as indicated in (**b**).

**i-j**, Heatmaps showing the distribution of log_2_ fold differences of FPKM normalized H3K27me3 signal at the gene body and surrounding loci of 184 genes significantly affected upon STM treatment. Genes were ordered by expression difference as indicated in (**b**).

**k**, Heatmaps showing the mean log_2_ fold difference in mean FPKM normalized H3K27me3 levels between STM2457 stimulated and control cells at 2,943 enhancers identified in^29^ and the +/-2 kbp surrounding regions.

**l-q**, Heatmaps showing the distribution of FPKM normalized H3K27me3 signal at 2,943 enhancers identified in^29^ and their surrounding loci. Enhancers were ordered by the mean difference in H3K27me3 signal as indicated in (**k**).

**r-s**, Heatmaps showing the distribution of log_2_ fold differences of FPKM normalized H3K27me3 signal at 2,943 enhancers identified in^29^ and their surrounding loci. Enhancers were ordered by the mean difference in H3K27me3 signal as indicated in (**k**).

**Supplementary Figure 5. Depletion of m^6^A Promotes *Eras* Dependent Germline Entry Deficiency in Mouse FS cells.**

**a-b**, *Oct4*ΔPE::GFP reporter mESCs were transduced either with scramble control (Scr), *Mettl3* (shM3-1) or *Mettl14* (shM14-1) shRNA, followed by Formative Stem (FS) cell induction.

**a**, Schemata illustrating experimental strategies. ActA_low_, low concentration of Activin A. XAV, WNT agonist XAV939. BMS, Retinoic Acid Receptor inhibitor BMS493. See also Figure S2m for other abbreviations.

**b**, qPCR showing *Eras* expression at day 2 of FS cell induction. Mean ± SD. N =4; two-tailed, paired t-test: vs each Scr, **p <0.01; * p<0.05.

**c**, ATAC-seq from a published dataset^2^ showing open chromatin peaks on the *Oct4* distal enhancer (DE) and proximal enhancer (PE) loci in mESCs versus FS cells. See Method for details.

**d-i**, Day 10 FS cells induced from *Blimp1*-GFP reporter ESCs, were transduced with scramble control (Scr), *Mettl3* (shM3-1), *Mettl14* (shM14-1) or *Mettl3* and *Eras* (shM3-1 sh*Eras*-2) shRNA for a further 10 days. These FS were either (**e-f, h**) collected for immediate analysis, or (**g**) directly induced into PGCLC, or (**i**) treated with DMSO or PI3K inhibitor LY294002 before PGCLC specification. See also Methods for experiment details. Mean ± SD. Two-tailed, paired t-test: vs Scr, \*\**p* <0.01; * *p*<0.05; vs shM3-1, ^##^*p* <0.01; ^#^*p* <0.05; N.S., Non-significant.

**d**, Schemata showing experimental strategies for shRNA knock-down and inhibitor treatments in self-renewing formative stem (FS) cells. ROCKi, ROCK inhibitor Y-27632; PVA, Poly (vinyl alcohol). See also (**a**) and Figure S2m for other abbreviations.

**e**, Colorimetric m^6^A quantification analysis in FS cells. N =3.

**f**, Morphology of *Blimp1*-GFP FS cells. Scale bar, 100 μm.

**g**, qPCR showing the expression of germline and pluripotency markers in day 4 PGCLC spheres. N =4.

**h**, qPCR showing *Fgf4* and *Fgf5* expression in FS cells. N =3.

**i**, Graphs summarizing FACS analyses measuring the percentage of SSEA1 and CD61 double positive cells in day 4 PGCLC spheres. N =3. See Supplementary Files for representative FACS plots.

**j-l**, *Blimp1*-GFP reporter FS cells were treated with DMSO control (Ctr) or METTL3 inhibitor STM2457 (STM) together with or without LY294001 treatment for 2 days. Mean ± SD. N =3; two-tailed, paired t-test: vs Ctr, \*\**p* <0.01; \**p* <0.05; vs STM,^#^*p* <0.05.

**j**, Colorimetric m^6^A quantification in FS cells.

**k**, Immunoblots of ERAS protein levels in FS cells.

**l**, Graph summarizing FACS analyses in day 4 PGCLC spheres showing percentage of SSEA1 and CD61 double positive cells. See Supplementary Files for representative FACS plots.

**m-o**, *Blimp1*-GFP reporter FS cells were treated with 4 days of DMSO (Ctr), 2 days of DMSO and 2 days of METTL3 inhibitor STM2457 (STM) or 2 days of STM2457 treatment followed by 2 days of STM2457 withdrawal (STMwd). Thereafter, PGCLC induction commenced.

**m**, Schemata illustrating experimental strategy.

**n**, Immunoblots of ERAS protein levels in *Blimp1*-GFP FS cells.

**o**, Graph summarizing FACS analysis showing the proportion of SSEA1 and CD61 double positive cells in day 4 PGCLC spheres. Mean ± SD. N =4; two-tailed, paired t-test: vs Ctr, \*\**p* <0.01; vs STM, ^##^*p* <0.01. See Supplementary Files for representative FACS plots.

**p**, Immunoblots showing ERAS level and AKT phosphorylation in R1 FS cells transduced with the over-expressing control lentivirus (OE Ctr) or Eras-overexpressing lentivirus (OE Eras).

**q-r**, Immunoblots showing YTHDF1, YTHDF3 and ERAS protein levels (**q**) and AKT phosphorylation (**r**) in *Blimp1*-GFP FS cells transfected with negative control (Nc) or *Ythdf1* (siY1) or *Ythdf3* (siY3) siRNA for 2 days.

**Supplementary Figure 6. Depletion of m^6^A Promotes Mesodermal Differentiation in Mouse FS Cells.**

**a**, Graph summarizing FACS analyses showing the percentage of *Blimp1*-GFP reporter positive cells in FS cells upon DMSO control (Ctr) or METTL3 inhibitor STM2457 (STM) treatment for 2 days. Mean ± SD. N =4; two-tailed, paired t-test: vs 0, \**p* <0.05; \*\**p* <0.01; vs 2, ^##^*p* <0.01. See also Supplementary Files for representative FACS plots.

**b**, Graph summarizing FACS analyses showing the proportion of *Blimp1*-GFP reporter positive FS cells. Cells were analyzed after treatment with DMSO for 4 days (Ctr), 2 days of DMSO followed by 2 days of METTL3 inhibitor STM2457 (STM) or 2 days of STM2457 treatment followed by 2 days of STM2457 withdrawal (STMwd). Mean ± SD. N =3; two-tailed, paired t-test: vs Ctr, \**p* <0.05; vs STM, ^##^*p* <0.01; N.S., Non-significant. See also Figure S5m for experimental design.

**c-d**, Immunoblots showing BLIMP1 protein levels in *Blimp1*-GFP reporter FS cells. For (**c**), FS cells were transduced with scramble control (Scr), *Mettl3* (shM3-1) or *Mettl3* and *Eras* (shM3-1 sh*Eras*-1 or shM3-1 sh*Eras*-2) shRNA. For (**d**), FS cells were transduced with scramble control (Scr) or *Eras* (sh*Eras*-1 or sh*Eras*-2) shRNA, followed by DMSO control or METTL3 inhibitor STM2457 (STM) treatment. See also Figure S5d for experimental design.

**e-g**, Graph summarizing FACS analyses showing the percentage of *Blimp1*-GFP reporter positive cells in the FS condition. For (**e**), FS cells were transduced with scramble control (Scr) or *Eras* (sh*Eras*-2) shRNA, followed by DMSO control (Ctr) or METTL3 inhibitor STM2457 (STM) treatment. For (**f**), FS cells were treated with DMSO control, METTL3 inhibitor STM2457 (STM) or STM2457 and PI3K inhibitor LY294002 (STM LY) for 2 days. For (**g**), FS cells were transduced with scramble control (Scr), *Mettl3* (shM3-1) or *Mettl14* (shM14-1) shRNA, followed by DMSO control or PI3K inhibitor LY294002 (LY) treatment. Mean ± SD. N ≥3; two-tailed, paired t-test; vs Scr or Ctr, \*\**p* <0.01; \**p* <0.05; vs shM3-1, ^##^*p* <0.01; vs Scr STM or STM, ^#^*p* <0.05. See also Figure S5d for experimental design and Supplementary Files for representative FACS plots.

**h**, Diagram showing the putative m^6^A motif sites on *Blimp1* mRNA and the primers designed for meRIP-qPCR. UTR, untranslated region. CDS, coding sequence.

**i-k**, *Blimp1*-GFP reporter FS cells were transduced with scramble control (Scr) or *Mettl3* (shM3-1) shRNA, then sorted into GFP positive and negative cell fractions. See also Methods for experimental details.

**i**, qPCR of *Blimp1* mRNA levels in *Blimp1*-GFP reporter positive and negative sorted cell fractions. Mean ± SD. N =3; two-tailed, paired t-test: vs Scr GFP -, \**p* <0.05; vs shM3-1 GFP -, ^#^*p* <0.05; N.S., Non-significant.

**j**, FACS analysis showing *Blimp1-*GFP fluorescence sorted population enrichment. SSC, side scatter parameter.

**k**, Immunoblots of protein levels for GFP, METTL3, EOMES, ERAS and pAKT in *Blimp1*-GFP reporter positive and negative sorted FS cells.

**l**, FACS analysis of *Blimp1*-GFP reporter positive and negative gated cells showing the percentage of FLK1 co-expression in FS cells, treated with DMSO control (Ctr) or METTL3 inhibitor STM2457 (STM) for 2 days.

**m**, Graph summarizing FACS data (**l**). Mean ± SD. N =3; two-tailed, paired t-test: vs Ctr GFP -, \**p* <0.05; vs STM GFP -, ^#^*p* <0.05; N.S., Non-significant.

**n**, Graph summarizing FACS analyses of *Blimp1*-GFP reporter positive and negative gated FS cell populations, showing the percentage of FLK1 co-expression. FS cells were transduced with scramble control (Scr) or *Mettl3* (shM3-1) shRNA. Mean ± SD. N =4; two-tailed, paired t-test: vs Scr GFP -, \*\**p* <0.01; vs shM3-1 GFP -, ^##^*p* <0.01; N.S., Non-significant. See also Supplementary Files for representative FACS plots.

**o**, qPCR showing *Flk1* expression in day 4 PGCLC spheres derived from *Blimp1*-GFP reporter FS cells, transduced with scramble control (Scr), *Mettl3* (shM3-1) or *Mettl3* and *Eras* (shM3-1 sh*Eras*-2) shRNA. Mean ± SD. N =4; two-tailed, paired t-test: vs Scr, \**p* <0.05; vs shM3-1, ^##^*p* <0.01; N.S., Non-significant.

**p**, Graph summarizing FACS analyses showing the percentage FLK1 positive cells in day 4 PGCLC spheres. Spheres were derived from *Blimp1*-GFP reporter FS cells, transduced with scramble control (Scr) or *Mettl3* (shM3-1) shRNA followed by DMSO control or PI3K inhibitor LY294002 (LY) treatment for 2 days before PGCLC induction. Mean ± SD. N =3; two-tailed, paired t-test: vs Scr, \**p* <0.05; vs shM3-1, ^#^*p* <0.05; N.S., Non-significant. See also Figure S5d for experimental design and Supplementary Files for representative FACS plots.

**q**, Immunoblots showing protein levels of mesodermal (MESP1 and MIXL1) and endodermal (FOXA2 and GATA4) markers in day 4 PGCLC spheres. Spheres were derived from *Blimp1*-GFP reporter FS cells, transduced with scramble control (Scr), Mettl3 (shM3-1) or *Mettl3* and *Eras* (shM3-1 sh*Eras*-2) shRNA.

**r**, Graph summarizing FACS analyses in Figure 6j showing the percentage of FLK1 positive and E-Cadherin negative mesodermal cells. Cells analysed for mesodermal lineage commitment were derived from *Blimp1*-GFP reporter FS cells, transduced with scramble control (Scr), *Mettl3* (shM3-1) or *Mettl3* and *Eras* (shM3-1 sh*Eras*-2) shRNA. Mean ± SD. N =3; two-tailed, paired t-test: vs Scr, \**p* <0.05; vs shM3-1,^#^*p* <0.05; N.S., Non-significant.

**Supplementary Figure 7. Depletion of m^6^A Impedes Human Germline Entry and Promotes Neuroectoderm.**

**a**, FACS analysis showing the percentage of *OTX2*-YFP reporter positive cells in DMSO control or METTL3 inhibitor STM2457 treated *OTX2*-YFP 4i hPSCs. Dashed line indicates positive gating. YFP signals from wild-type H1 4i hPSCs served as a negative control. See also Figure 7e for experimental repeats.

**b**, qPCR analyses showing the expression of pluripotency and lineage markers in H9 4i hPSCs treated with DMSO (Ctr) or METTL3 inhibitor STM2457 (STM) for 3 days. Mean ± SD. N =3; two-tailed, paired t-test: N.S., Non-significant.

**c**, qPCR showing mRNA degradation rates after Actinomycin-D treatment in H9 4i hPSCs treated with DMSO control or METTL3 inhibitor STM2457 for 3 days. Mean ± SD. N =3; two-tailed, paired t-test: vs Ctr, \*\**p* <0.01; \**p* <0.05; N.S., Non-significant.

**d**, Immunoblots showing the phosphorylation of FGFR1, ERK and AKT in H9 4i hPSCs cultured on Mouse Embryonic Fibroblast (MEF) feeders or in MEF feeders alone. Cells were treated with DMSO control, METTL3 inhibitor STM2457 or STM2457 and FGFR1 inhibitor PD173074 or MEK inhibitor PD325901 for 3 days.

**e**, H9 4i hPSCs were treated with DMSO (Ctr), METTL3 inhibitor STM2457 (STM), or STM2457 and FGFR1 inhibitor PD173074, PI3K inhibitor LY294002 or MEK inhibitor PD0325901, followed by protein extraction. Immunoblots showing OTX2 levels.

**f**, Immunoblots showing the phosphorylation of AKT upon DMSO control, METTL3 inhibitor STM2457 or STM2457 together with FGFR1 inhibitor PD173074 or PI3K inhibitor LY294002 treatments for 3 days in H9 4i hPSCs.

**g**, qPCR in H9 4i hPSCs showing mRNA changes for FGF and FGFR family members upon DMSO or METTL3 inhibitor STM2457 (STM) treatment for 3 days. Mean ± SD. N =3; two-tailed, paired t-test: vs Ctr, \*\**p* <0.01; \**p* <0.05. The expressional changes for the rest genes are non-significant.

**h**, FACS analysis showing co-expression of human germline marker CD49f and *OTX2*-YFP in day 4 human PGCLC spheres derived from *OTX2*-YFP reporter 4i hPSCs, which were treated with DMSO control or METTL3 inhibitor STM2457 for 3 days before PGCLC induction.

**i-j**, H9 4i hPSCs were treated with DMSO and negative control siRNA (Ctr Nc), STM2457 and negative control siRNA (STM Nc) or STM2457 and *OTX2* siRNA (STM si*OTX2*) for 3 days before PGCLC induction. (**i**) Immunoblots showing OTX2 expression in the 4i cells. (**j**) Representative FACS analysis showing the percentage of human germline marker EpCAM and CD49f double positive cells in day 4 PGCLC spheres.

**k-m**, qPCR showing (**k**) mesoderm, (**l**) neuroectoderm and (**m**) endoderm lineage marker expression levels. H9 4i PSCs were treated with DMSO control or METTL3 inhibitor STM2457 for 3 days, prior to differentiation into mesoderm, neuroectoderm or endoderm. Mean ± SD. N =3; two-tailed, paired t-test: vs Ctr, \*\**p* <0.01; \**p* <0.05; N.S., Non-significant. See also Methods for the experiment details.

**n**, Consensus m^6^A motifs identified in biological replicate 2 of R1 FS cells and H9 4i hPSCs in the DMSO control condition.

**o**, Radar plot of m^6^A peak distribution in R1 FS cells and H9 4i hPSCs in the DMSO control condition.

