## Supplemental Figures for "N^6^-Methyladenosine Safeguards Mouse and Human Germline Competence"

### Supplementary Figure 1

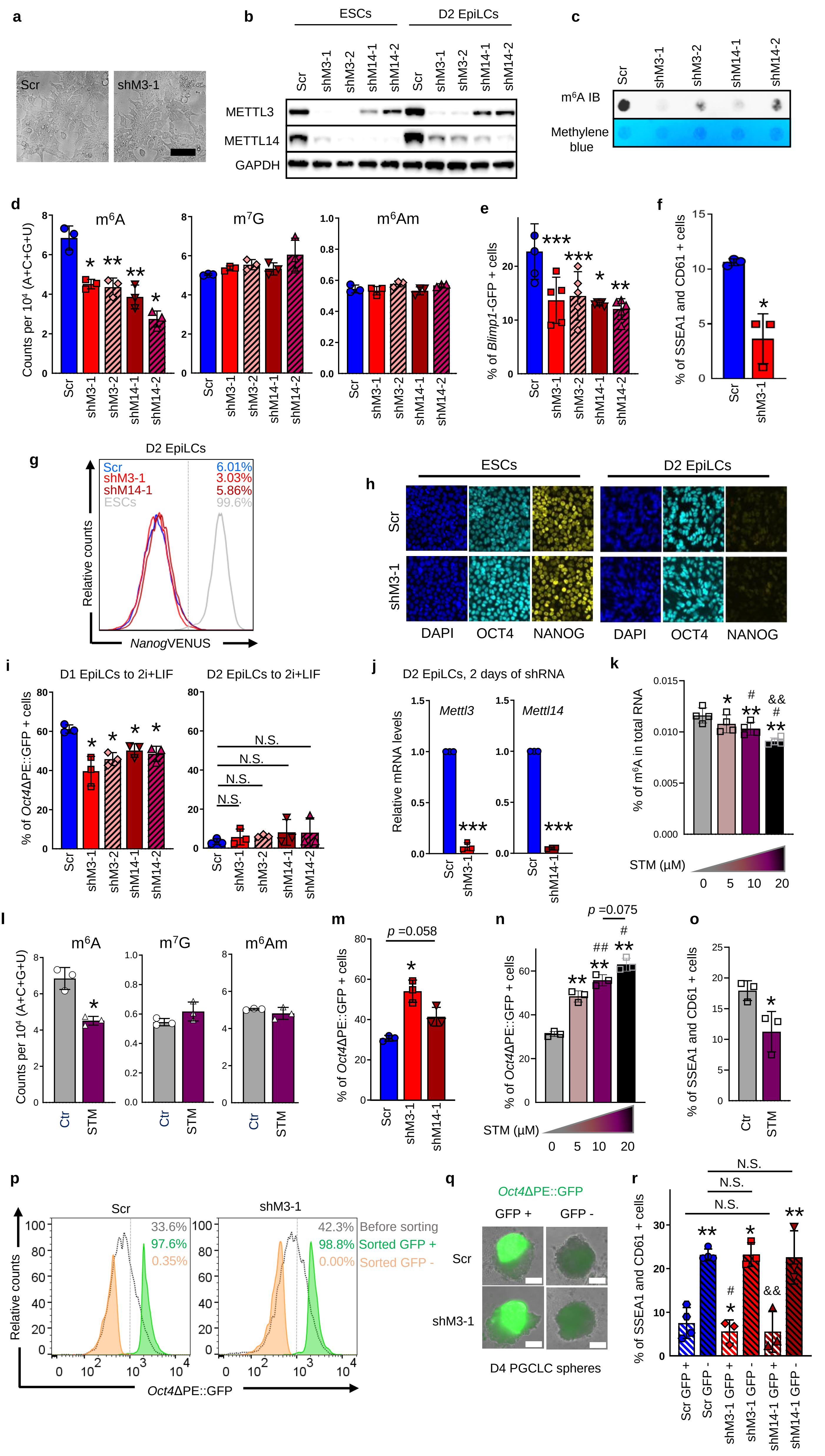

Supplementary Figure 2

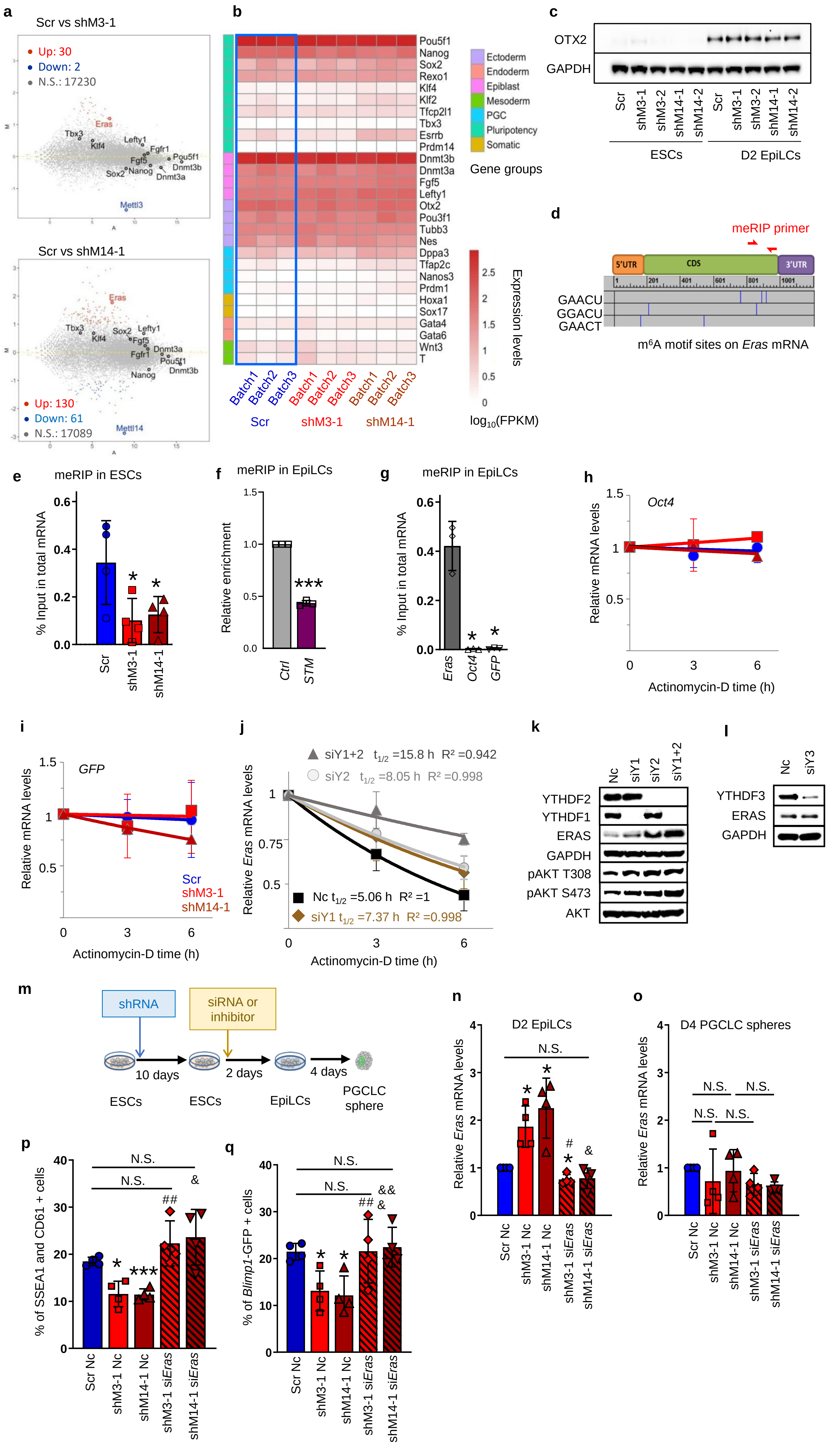

### Supplementary Figure 3

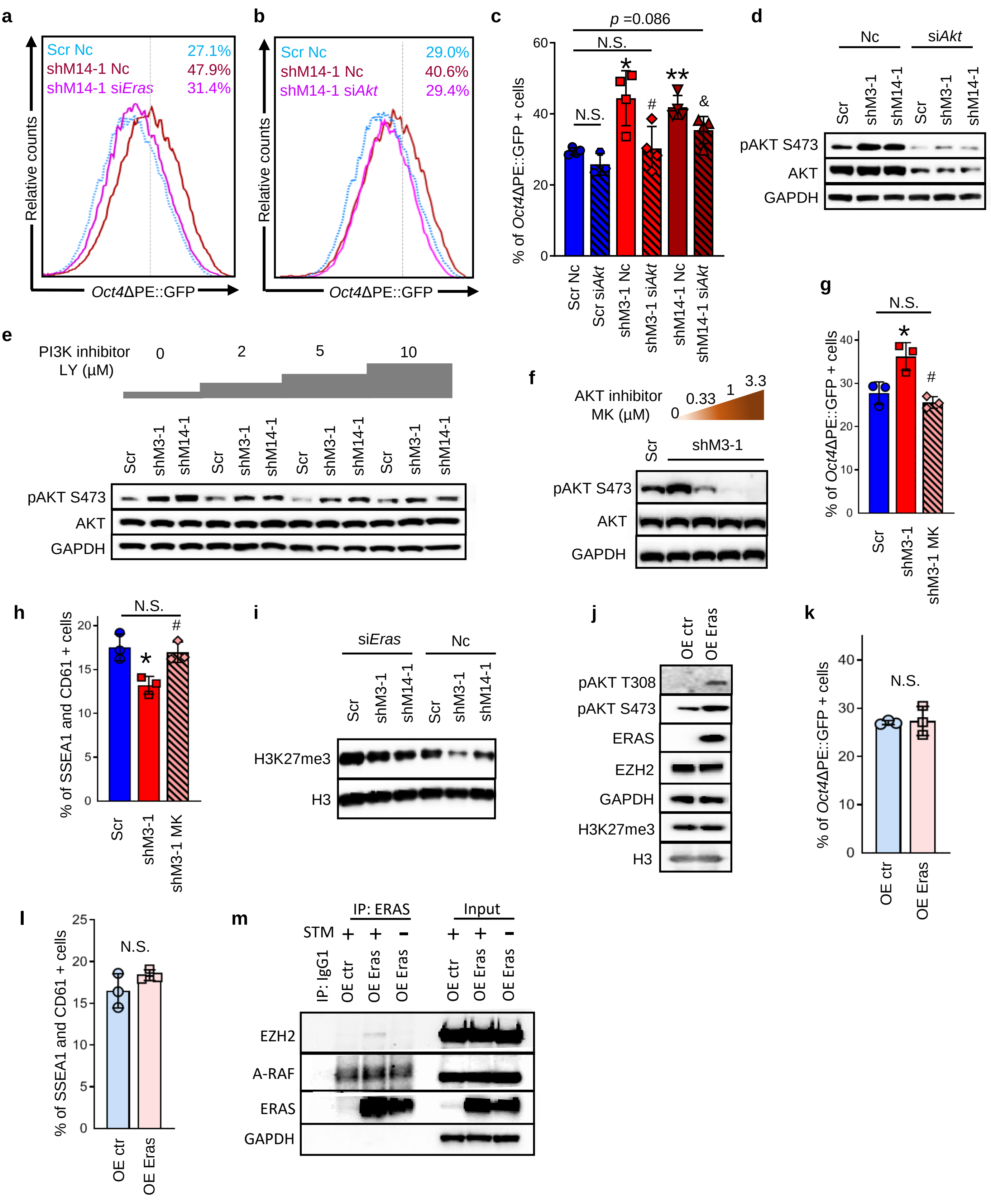

**Supplementary Figure 4**

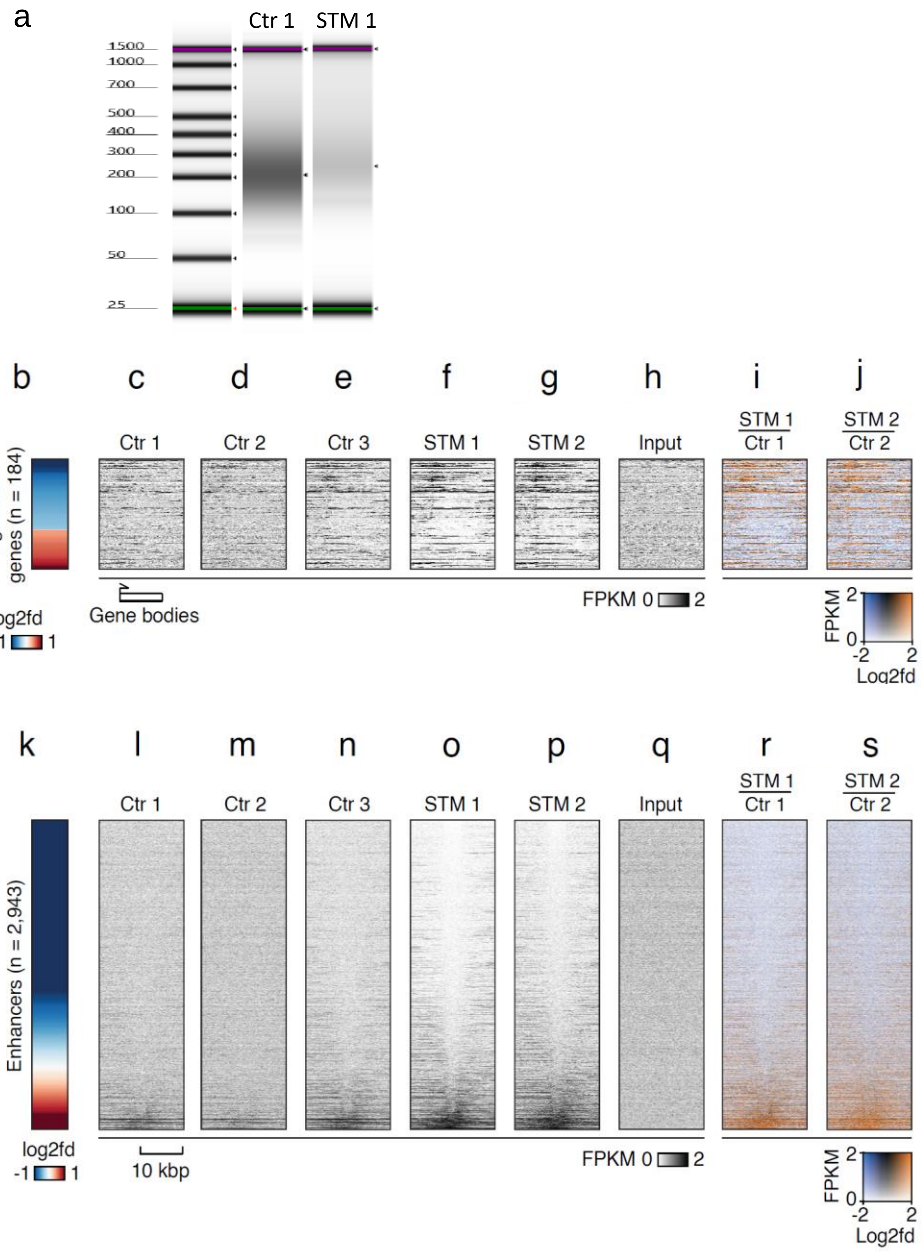

Supplementary Figure 5

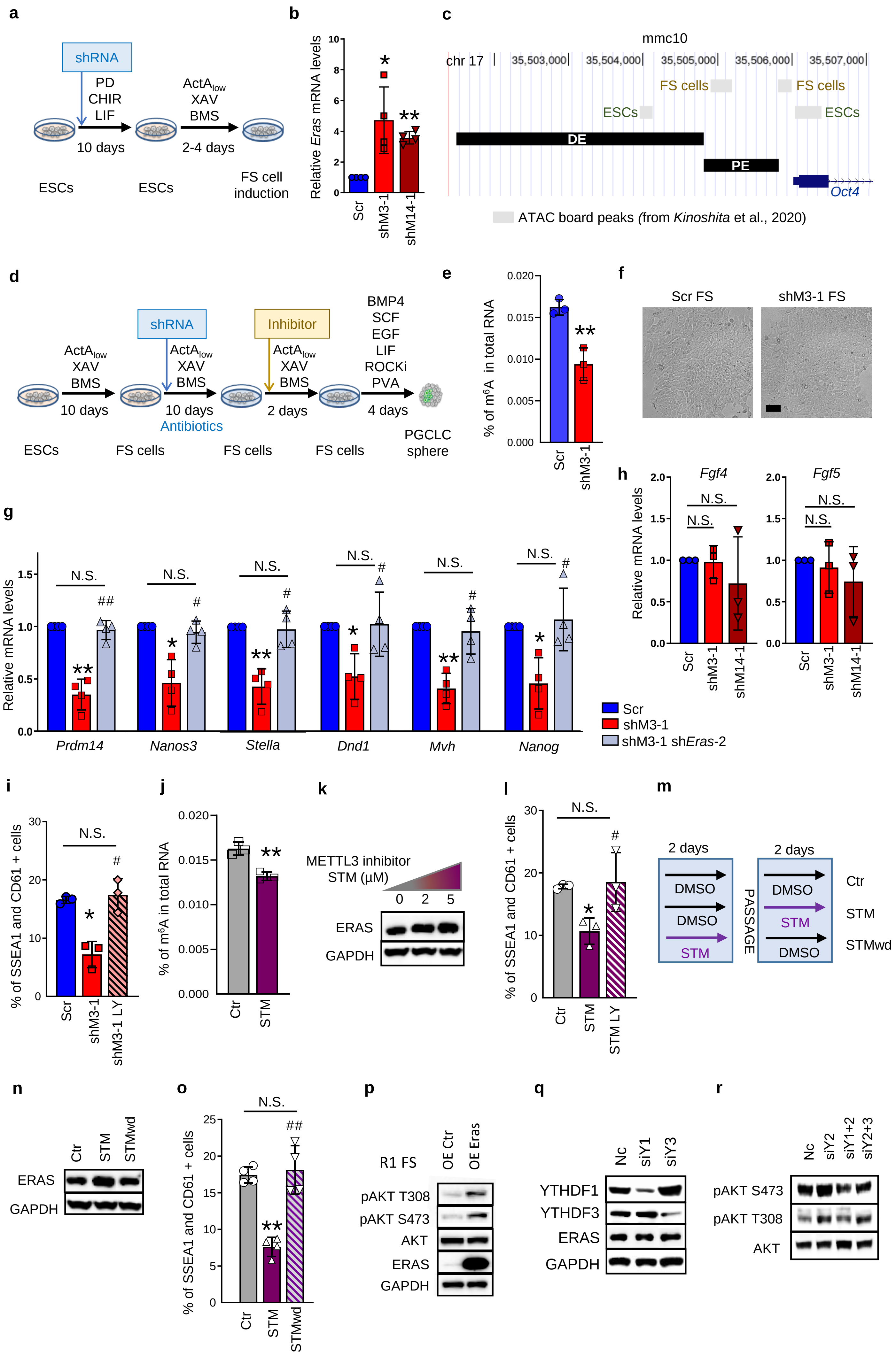


2025/11/14

**Supplementary Figure 6**

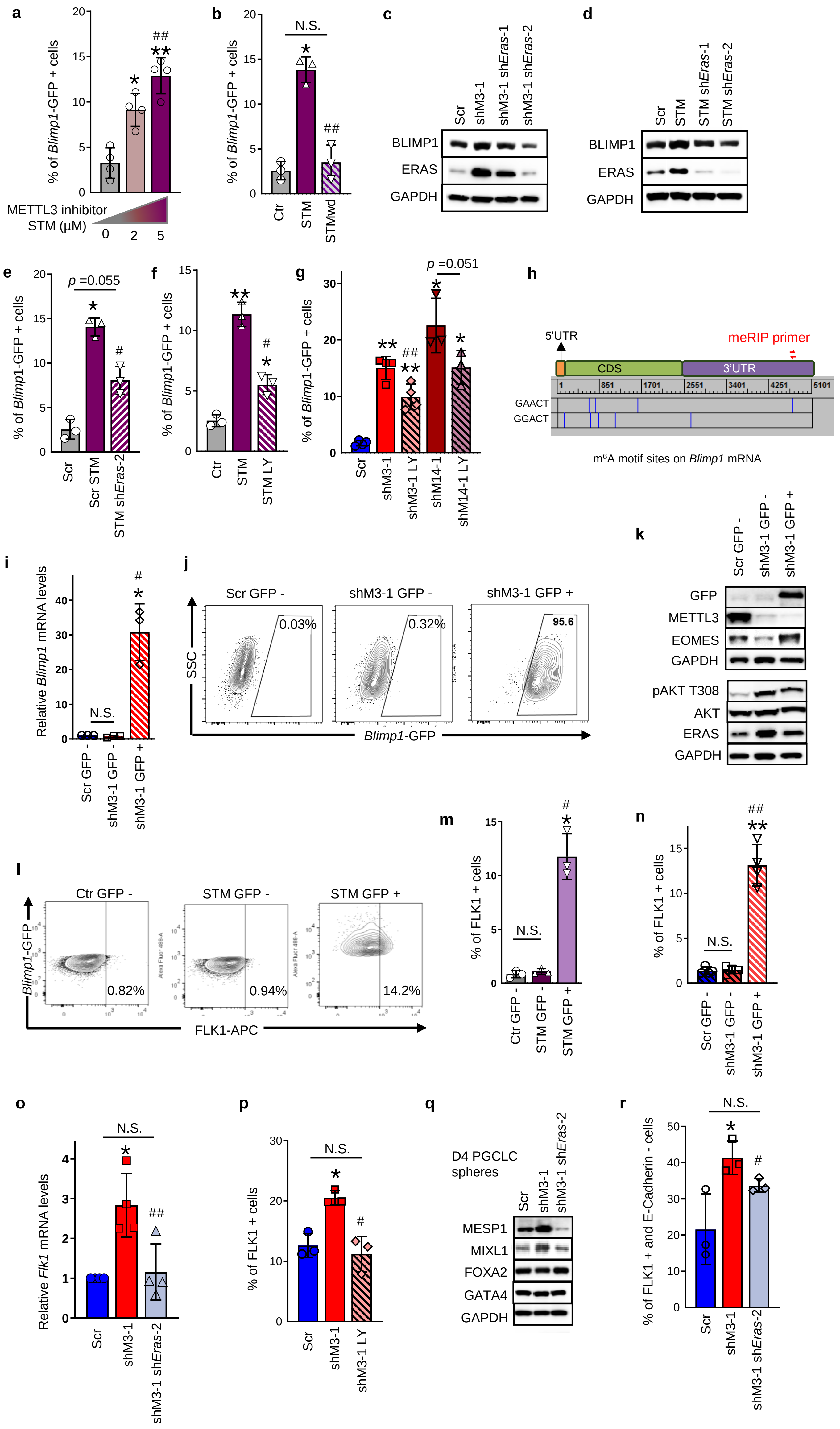

### Supplementary Figure 7

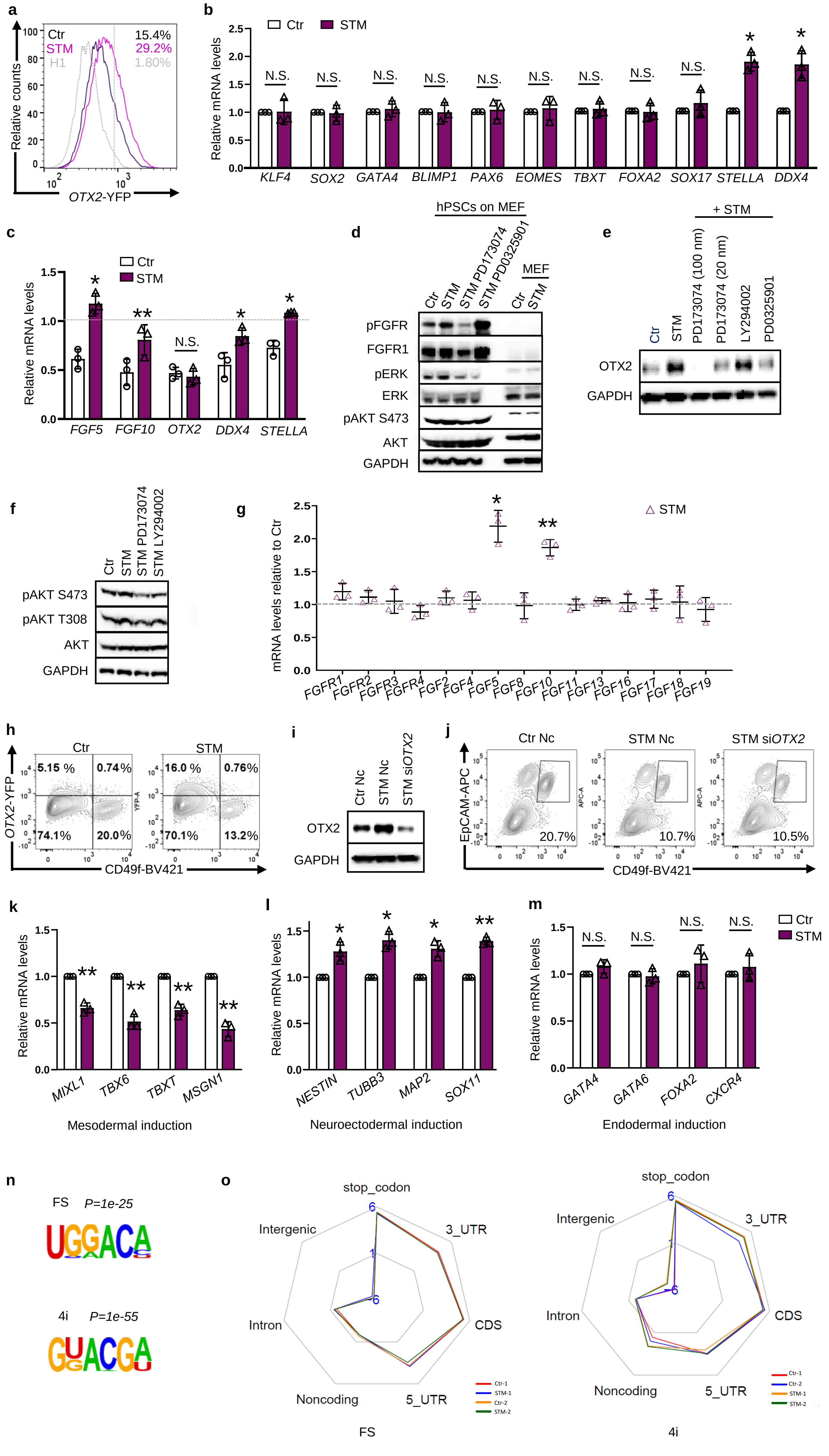
